# Reactive suppression of distractor representations resolves multidimensional interference

**DOI:** 10.1101/2025.11.16.688701

**Authors:** Davide Gheza, Michael C. Freund, Thea R. Zalabak, Wouter Kool

## Abstract

Navigating competing attentional demands is a core cognitive function, yet how the brain tunes control in multidimensional environments remains poorly understood. Here, we used a multidimensional task-set interference paradigm in which participants attended to one of four stimulus dimensions while three others acted as distractors, combining multivariate decoding, representational similarity analysis, and encoding models applied to human EEG. Targets and distractors were initially encoded in parallel, but distractor representations were rapidly suppressed ∼250 ms after stimulus onset, with suppression scaling with each distractor’s own conflict history. Neither trial-to-trial adaptation nor block-level learning produced anticipatory changes in task-relevant representations. Instead, proactive control modulated the speed and efficiency of stimulus-triggered suppression. Encoding models further revealed that conflict is represented in orthogonal, dimension-specific subspaces that eventually collapse onto a shared conflict signal. These results show that multidimensional attentional control operates through selective, reactive suppression of distractor representations, guided by a structured multivariate conflict signal.

## Introduction

Adaptive behavior requires the flexible integration of multiple sources of information, such as external stimuli, goals, and context. As you work through this text, you must hold your goal in mind, tune out background noise, and set aside whatever you were just thinking about. Our ability for attentional control^1^ enables such orchestration of processing, allowing us to occupy the brain’s *representational space*^2,3^ with attentional sets that emphasize goals and flexibly suppress transient distractors^4,5^. Contextual factors, such as prior knowledge of the text’s topic, can help establish appropriate control settings in an anticipatory, proactive way^6,7^. This raises a fundamental question: how does attentional control orchestrate multiple competing representations within a limited representational space?

Although daily life demands flexible switching between sources of information, traditional cognitive control research examines interference from a single distractor: a task-relevant feature signals the correct response while a second feature primes a congruent or an incongruent response^8–11^. A key insight from work in this domain is that interference adapts based on recent demands: after experiencing conflict, people become less sensitive to distractors^5^. A canonical framework proposes that the anterior cingulate cortex detects control demands and uses this signal to modulate the relative gain on task-relevant versus task-irrelevant representations (conflict monitoring theory)^4^, inspiring a generation of conflict monitoring theories^12–16^. Yet these models assert a single, global conflict signal, offering limited predictions for how attentional control operates in environments with multiple sources of distraction.

To address this gap, we recently developed the Multidimensional Task-Set Interference (MULTI) paradigm, in which participants attend to one of four stimulus dimensions (color, shape, edge, pattern) while the three non-cued dimensions act as simultaneous distractors^9,17^. Because task cues change every few trials, all distractors remain salient throughout the task^18,19^. Behavioral data revealed that control adaptation is *dimension-specific*: Conflict from one distractor dimension selectively reduces subsequent interference from that same dimension, without generalizing to others. This suggests that attentional control operates through selective modulation of dimension-specific representations, rather than a unitary global conflict signal, raising a direct question about the underlying neural mechanism.

These findings generate direct neural predictions. First, control should modulate distractor rather than target representations. Second, these modulations should be dimension-specific: previously incongruent distractors should be selectively suppressed, and previously congruent ones enhanced. Third, if adaptation is dimension-specific, conflict itself cannot be a scalar quantity: it must be encoded in structured, separable representational axes, a prediction that is inconsistent with classic conflict monitoring accounts. To test these predictions, we applied multivariate decoding and representational similarity analysis (RSA)^20^ to human EEG recordings and developed a complementary encoding model to probe the representational structure of the conflict signal.

The temporal resolution of EEG further allowed us to ask whether control adaptation reflects reactive or proactive control adjustments^6^: reactive control refers to within-trial adjustments triggered by stimulus onset; proactive control refers to sustained adjustments carried across trials based on prior experience, which pre-empt distractor capture. Although a few studies have provided evidence for reactive suppression^21–23^ (see also ^24,25^), these primarily characterized adaptation within rather than across trials, and for a single distractor dimension.

To probe sustained control learning, we also introduced a task-specific proportion congruency (TSPC) manipulation, so that each task dimension was stably associated with a high, neutral, or low probability of distractor interference across blocks^7^. This design allowed us to ask whether participants learn long-term associations between task dimensions and expected conflict levels, and whether any such learning is expressed proactively, through anticipatory changes in task representations, or reactively, upon stimulus onset.

To foreshadow the core findings, we found that neural distractor representations are selectively suppressed in proportion to their conflict history, in a dimension-specific fashion consistent with our behavioral and computational predictions. Critically, both trial-to-trial and block-level (TSPC) adaptations were expressed through the same reactive suppression mechanism, with proactive learning tuning its speed and efficiency rather than establishing anticipatory representational states. The encoding models revealed that conflict is represented in orthogonal, dimension-specific subspaces that only converge onto a shared monitoring signal late within the trial. Together, these results challenge global accounts of cognitive control regulation and suggest that the brain manages multidimensional interference through selective, reactive suppression of distractor representations, guided by a structured multivariate conflict signal.

## Results

### Multidimensional interference and conflict adaptation

We recorded EEG while twenty-seven participants performed the MULTI (Fig. 1A)^17^. On each trial, they viewed two objects that varied across four dimensions – color (blue/orange), shape (oval/rectangle), edge (solid/dashed), and pattern (horizontal/vertical) – and selected the object containing the target feature for the dimension that was cued before stimulus onset. Target features were randomly determined at the start of the experiment and remained constant throughout. The remaining three dimensions acted as distractors, each independently congruent (target feature on the same side as the cued target) or incongruent (on the opposite side; Fig. 1B), yielding a parametric congruency scale from 0 to 3. The cued dimension switched every few trials, keeping all dimensions periodically relevant and salient throughout the session (Fig. 1C).

**Figure 1.**
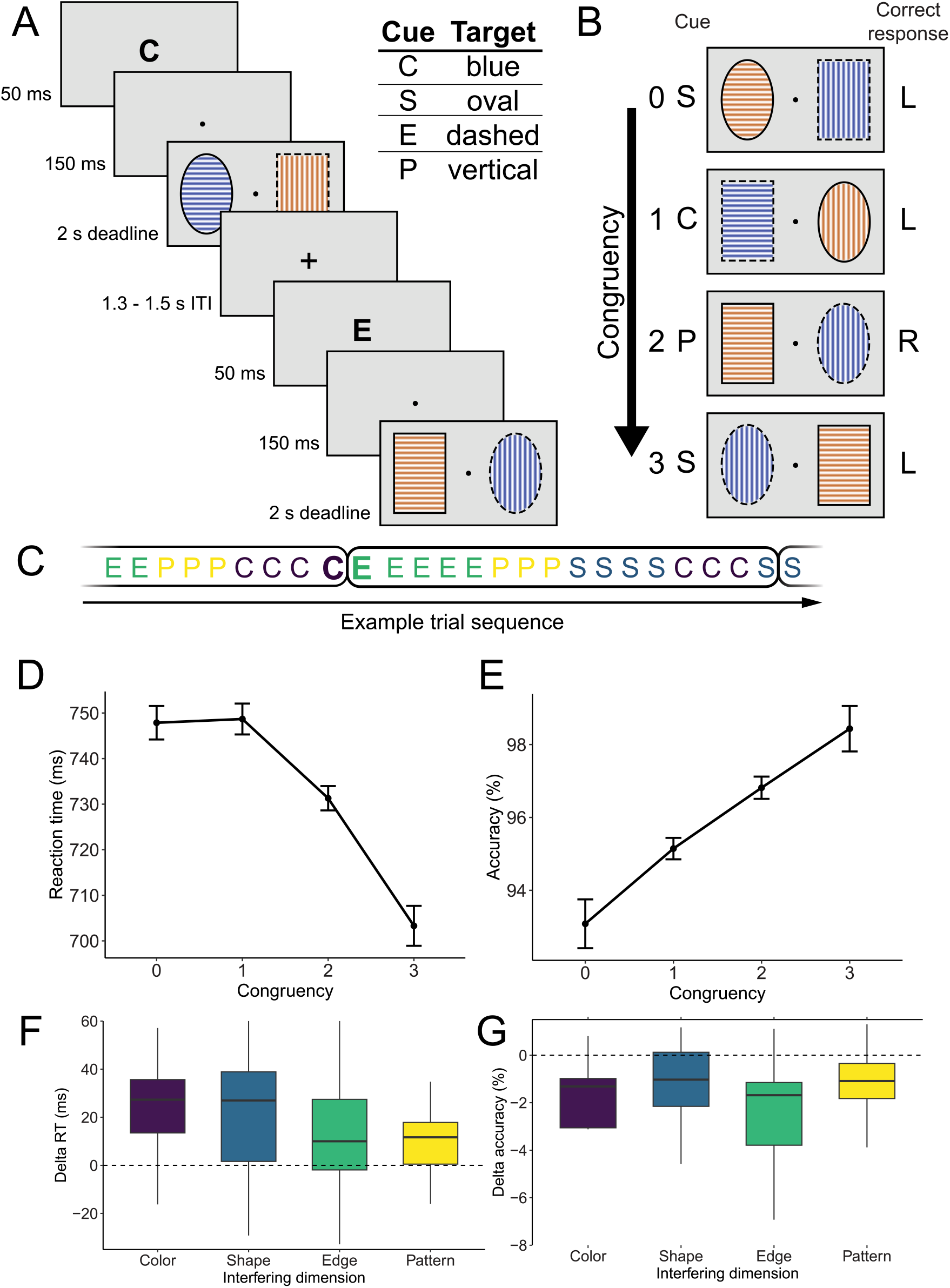
Multidimensional task-set interference and congruency effects (*N* = 26). **a**, Sequence of two example trials. On each trial, a letter cue (50 ms) indicated the task-relevant dimension, followed by a fixation dot (150 ms) and two stimuli presented side by side until response or deadline (2 s). Each stimulus was defined by four visual dimensions: colour (orange/blue), shape (oval/rectangle), edge (solid/dashed), and pattern (horizontal/vertical), with features randomly assigned to each stimulus on every trial. Participants selected the object containing the target feature for the cued dimension, each of which was instructed at the start of the task. **b**, Congruency was parametrically manipulated from 0 (all three non-cued dimensions prime the opposite side from the target) to 3 (all three prime the correct side). **c**, Example trial sequence. The cued dimension repeats for 3–5 trials before switching, with all four dimensions appearing within every 4 switches (encircled). **d, e**, Increasing congruency improved RT (**d**) and accuracy (**e**) monotonically. **f, g**, Interference effects of each distractor dimension (incongruent – congruent trials) on RT (**f**) and accuracy (**g**). All dimensions generate reliable interference, with the pattern dimension showing the weakest effect. Error bars represent within-subject 95% confidence intervals. Box centers, hinges, and whiskers indicate medians, interquartile ranges, and 1.5× interquartile range, respectively, and outliers are omitted.

Behavioral results, analyzed using generalized Bayesian linear regressions, confirmed that participants were sensitive to the cumulative interference from all non-cued dimensions (Fig. 1DE). Reaction times decreased and accuracy increased monotonically as more distractors supported the correct response (evidence ratio, ER = ∞ for the linear effect of congruency). Each non-cued dimension produced reliable interference (Fig. 1FG), defined as a performance difference between congruent and incongruent trials, for both response time (RT; color: Bayes Factor (BF) = 6.5 × 10^4^, shape: BF = 448, edge: BF = 6, pattern: BF = 24) and accuracy (color: BF = 766, shape: BF = 12, edge: BF = 614, pattern: BF = 70). Although all dimensions generated interference, there was substantial heterogeneity in their potency, with the pattern dimension exerting the weakest effect.

Attentional control adapted parametrically to recent conflict (Fig. 2AB). Following high-congruency trials, participants’ sensitivity to interference was enhanced, whereas following low-congruency trials, it was reduced. This pattern, consistent with a congruency sequence effect^5^ (CSE), was confirmed by strong evidence for an interaction between current and previous congruency on both RT and accuracy (ERs = ∞). These effects were substantial in magnitude. Relative to fully incongruent trials, fully congruent trials increased interference by 50 ms and -7.2% in RT and accuracy on following high-conflict trials, and increased facilitation effects by -36 ms and 1.2% on following low-conflict trials.

**Figure 2.**
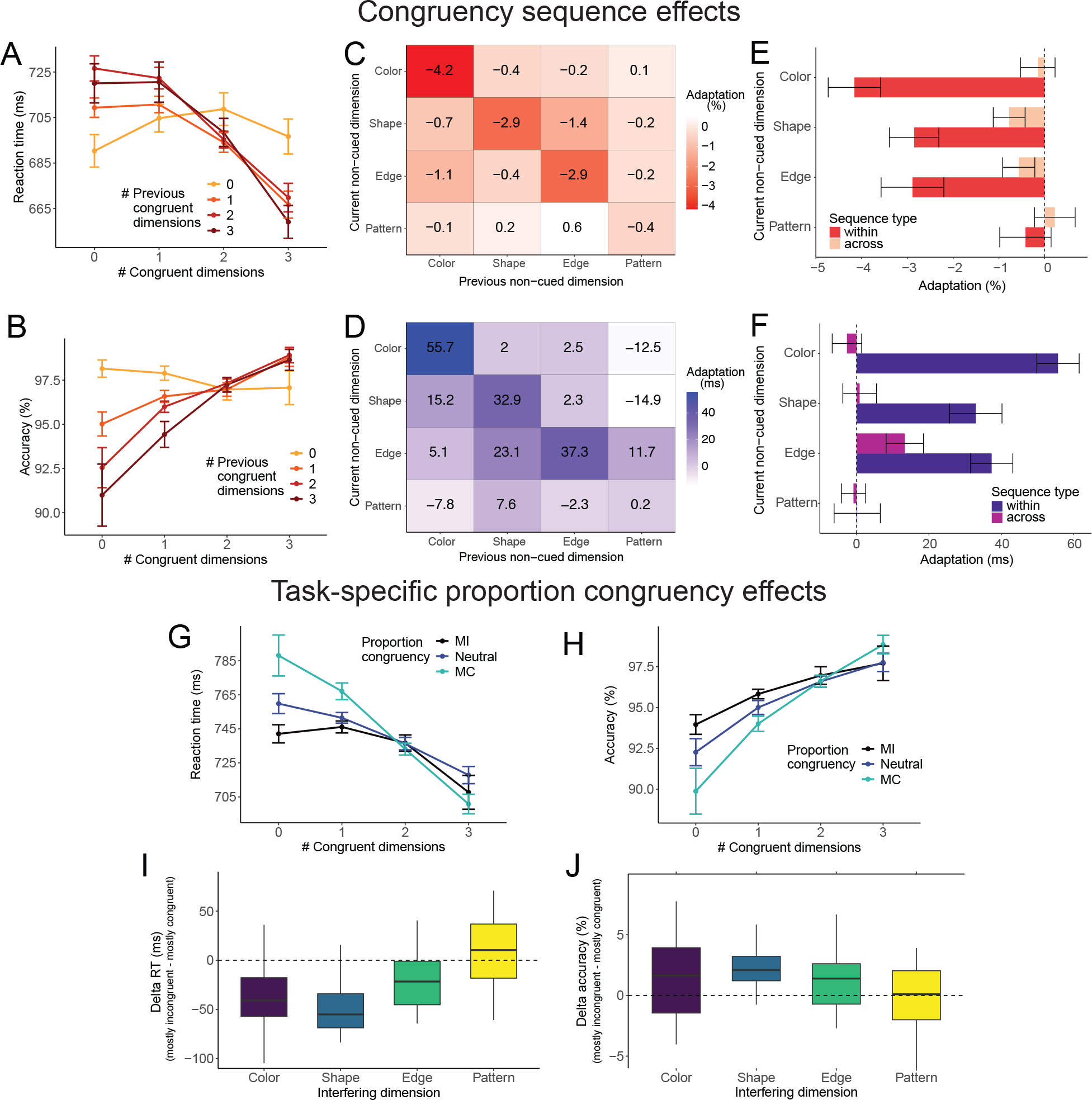
Behavioral congruency sequence (top) and task-specific proportion congruency effects (bottom). **a**,**b**, Parametric congruency sequence effects. Sensitivity to current congruency (x-axis) was modulated by the congruency on the preceding trial (color-coded): following low-congruency trials the congruency effect was attenuated, but following high-congruency trials it was enhanced, reflecting both stronger facilitation and interference under relaxed control. **c**,**d**, Heat maps of trial-to-trial conflict adaptation across all 16 combinations of previous and current non-cued dimensions for RT (**c**) and accuracy (**d**). Diagonal cells index within-dimension effects (e.g., the effect of previous color congruency on current color sensitivity), whereas off-diagonal cells index cross-dimension effects. Adaptation is defined as the change in the current dimension’s interference following congruent versus incongruent trials on the previous dimension. Effects are largely confined to the diagonal, indicating high dimension-specificity. **e**,**f**, Summary of within-dimension (diagonal; darker) versus across-dimension (off-diagonal; lighter) adaptation effects. Within-dimension adaptation was reliable for color, shape, and edge, but not pattern. **g**,**h**, TSPC effects on RT (**g**) and accuracy (**h**). Block-wise congruency context (mostly incongruent, MI, neutral, mostly congruent, MC) modulated sensitivity to current congruency, such that interference was reduced under MI and enhanced under MC contexts. **i**,**j**, TSPC-driven adaptation by interfering dimension for RT (**i**) and accuracy (**j**) computed as the difference in their interference effects between MI and MC blocks. Data are shown as means with within-subject 95% confidence intervals (**a, b, g, h**) or as boxplots showing median, interquartile range, and ±1.5× IQR whiskers (**e, f, i, j**).

To test whether these adjustments generalize across multidimensional distractors, we compared adaptation within versus across non-cued dimensions, asking whether conflict from a given dimension modulated subsequent interference from that same dimension (e.g., previous color congruency on current color sensitivity) or from others (e.g., previous color congruency on current shape sensitivity). Adaptation was strongly dimension-specific. Conflict from a given distractor reduced its own subsequent interference but did not transfer to other dimensions (Fig 2CD, diagonal; Fig. 2EF, darker colors). Within-dimension adaptation was robust for color (RT: BF = 3249005.3, accuracy: BF = 1540.1), shape (RT: BF = 287.1, accuracy: BF = 10.7), edge (RT: BF = 203.3, accuracy: BF = 20.6), but not for pattern (RT: BF = 0.2, accuracy: BF = 0.7), mirroring that dimension’s weaker interference. Across-dimension adaptation was consistently undetected (RT: range BFs = 0.2 - 1.9, accuracy: range BFs = 0.2 - 1.3; Fig. 2CD, off-diagonal; Fig. 2EF, lighter colors). These results, which replicate our previous findings ^17^, demonstrate that multidimensional control is guided by dimension-specific rather than global conflict signals.

### A computational model of dimension-specific adaptation

To characterize the within-trial temporal dynamics underlying these effects, we implemented a dynamic neural network model of the MULTI (Fig. 3B), extending the static model introduced in ^17^. This model combines evidence from four parallel dimension-specific pathways, each integrating stimulus input and top-down task signal via leaky accumulation. This architecture departed from prior models that measure conflict at the response level. Instead, conflict within each non-cued pathway is quantified as Hopfield energy^26^, a measure of the degree to which pathway activity is inconsistent with the correct response. This signal is then used to update pathway-specific control signals across trials. The model’s parameters were tuned to qualitatively capture the key behavioral signatures: parametric congruency effects, dimension-specific CSEs, and the relative weakness of the pattern dimension across both interference and adaptation measures.

**Figure 3.**
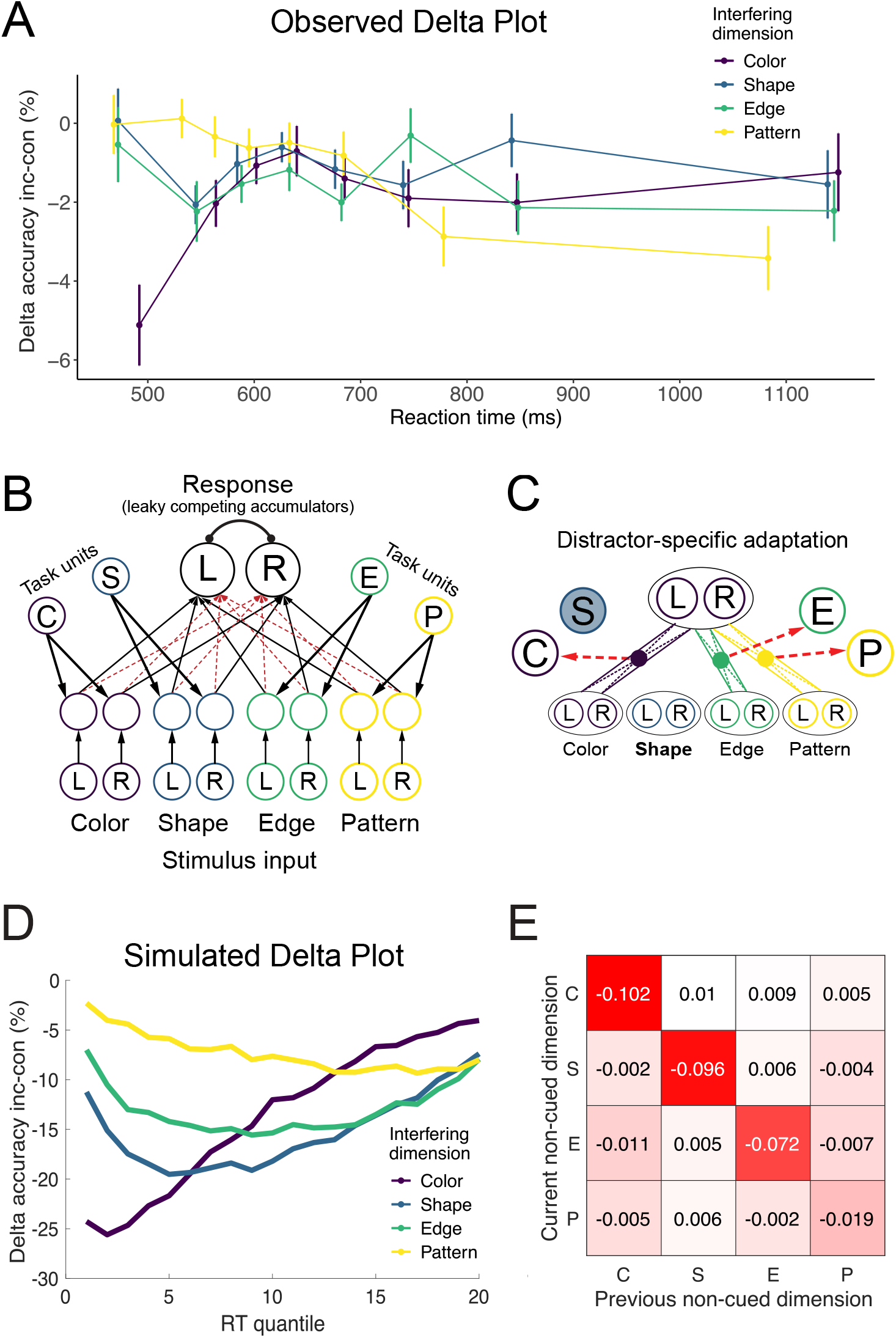
Model architecture and processing dynamics. **a**, Behavioral delta plots showing interference effects on accuracy across RT quantiles, split by dimension. Color interference emerges early, present even for the fastest responses, while pattern interference is absent for fast RTs and increases only for slower responses. **b**, Schematic of the dynamic LCA model. Stimulus and task inputs project onto four dimension-specific pathways, each integrating stimulus and task input via leaky competing accumulation with dimension-specific time constants (τ_d_). Hidden activity feeds into two response units that have balanced excitatory and inhibitory connections. Choice is determined by a pair of opposing noisy accumulators racing to threshold to yield accuracy and RT. Pathway-specific conflict is computed as Hopfield energy between hidden and response activity and used to modulate task-cue input on subsequent trials. **c**, Dimension-specific conflict-control loop. Conflict is computed as Hopfield energy within each non-cued dimension’s pathway, and transformed into a dimension-specific control value that modulates the corresponding task unit on the subsequent trial. This mechanism produces distractor-specific adaptation without cross-dimensional generalization. **d**, Simulated delta plots reproducing the heterogeneous temporal dynamics observed behaviorally (Fig. 2G), with color emerging early, and pattern interference affecting only slower responses. **e**, Simulated adaptation heatmap showing dimension-specific conflict adaptation across all 16 combinations of previous and current non-cued dimensions. Effects are confined to the diagonal, replicating the behavioral pattern (Fig. 2CD), and are weaker for the pattern dimension, consistent with its slower integration dynamics.

Our prior implementation treated conflict as stable within a trial. However, recent work suggests that interference effects and adaptation may unfold at a sub-trial timescale^21,27,28^. To test this, we used delta plots^29^, which track how interference effects on accuracy evolve across reaction time quantiles (Fig. 3A). Whereas prior delta-plot work has typically isolated single sources of conflict, the MULTI enables a within-task comparison of multiple dimensions. This analysis revealed heterogeneous temporal dynamics: with early interference for color and a gradual ramping of interference for the others. Combined with the estimated potency for each distractor (Fig.1FG), this suggests that a dimension’s interference potency may be governed by the speed of its visual integration.

To model these temporal dynamics of interference within a trial, we developed a dynamic leaky competing accumulator (LCA) variant of this model (Fig. 3B). In this model, each pathway integrates evidence according to its own temporal constant (τ), while mutual inhibition between accumulators yields competition between dimensions. This extension allows the model to reproduce the observed heterogeneity in the temporal evolution and strength of distractor interference, including the weaker and later emergence of the pattern dimension.

Simulations revealed that adaptation arises from selective suppression of distractor pathways rather than target enhancement (Fig. 3C). Dimension-specific adaptation emerged despite heterogeneity in the timing and strength of interference (Fig. 3E). Importantly, weaker dimensions indeed led to reduced adaptation (Fig. 3D), as we observed in our behavioral data (Fig. 2CE).

Together, the behavioral and modeling results converge on a clear neural prediction: adaptation should appear as dimension-specific modulations of the distractor representation strength, with the timing and strength of this adaptation scaling with each dimension’s interference potency. While frontal midline theta power replicated canonical signatures of conflict detection^30–33^ in the MULTI (see Fig. S1), univariate EEG measures are not well-suited to resolve the dimension-specific structure our predictions demand. Therefore, we turned our analytical framework to multivariate decoding of EEG, which allowed us to track the strength and temporal evolution of all four dimension-specific representations on a millisecond timescale.

### Task representations are encoded in parallel but unaffected by conflict history

To do this, we adopted the decoding-RSA framework developed by Kikumoto and Mayr^20^, which combines multivariate decoding with representational similarity analyses to track the strength and temporal evolution of task-related information (Fig. 4A). After successfully training a decoder to detect the eight possible cue–side conjunctions from EEG spectral power (Fig. 4B) at each timepoint in the trial, we used the resulting classification probability profiles as the substrate for RSA. Specifically, the decoder outputs a probability distribution across all eight possible cue-side conjunctions for each timepoint. This distribution does not only assign mass to the correct conjunction, but it also spreads probability across other dimensions. Importantly, this pattern of “confusions” carries information unrelated to the correct conjunction. By applying RSA model matrices to these distributions, we can estimate the strength of three task representations: the cued task rule, the spatial location of the target feature, and their conjunction, which captures feature-specific attention.

**Figure 4.**
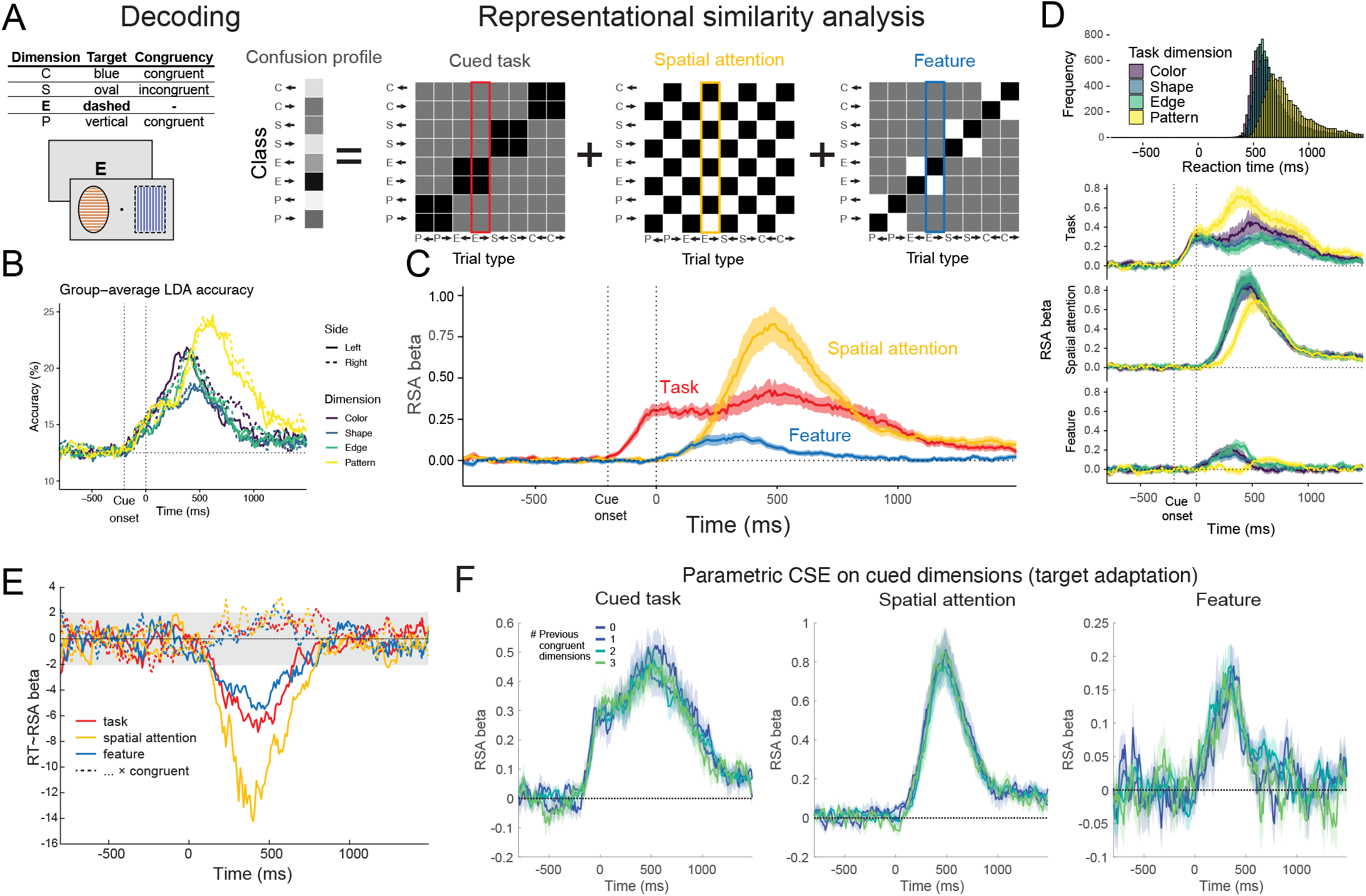
Decoding-RSA of task-relevant neural representations. **a**, Schematic of the single-trial decoding-RSA procedure. The classifier outputs a probability distribution across all eight cue-side conjunctions (i.e., confusion profile), which is regressed against columns from three RSA model to estimate the strength of the cued task rule, the location of spatial attention (an axis shared across dimensions), and their conjunction for each time step in each trial. **b**, Time-resolved decoding accuracy for the classifier trained to discriminate cue identity and target location from EEG spectral power, stratified by class. Dashed lines indicate chance. **c**, Temporal profiles of cued task, spatial attention, and feature coding strengths, averaged across dimensions. Shaded areas indicate ±1 SEM; vertical dashed lines mark the cue and stimuli onsets. **d**, RT distributions by dimension (top) and dimension-specific coding profiles (bottom). Spatial attention and feature coding are delayed for the pattern dimension, consistent with slower visual integration. **e**, Multi-level regression coefficients showing that stronger coding of all three representations predicts faster RTs (solid lines); the behavioral benefit was modestly reduced on high-congruency trials (dashed lines). **f**, RSA coding strength stratified by the number of congruent dimensions on the preceding trial, showing no evidence of modulation by prior conflict.

The temporal profile of decoded representations followed the structure of the trial (Fig. 4C). Cue coding emerged rapidly after cue onset and showed a bimodal profile, with an early plateau around stimulus onset followed by a second rise around 500 ms (plausibly reflecting its later integration with task-rule representations). Spatial attention coding, reflecting spatial orienting and response generation, emerged after stimulus onset and peaked just before the median reaction time (∼500 ms). Their conjunction, capturing feature-specific spatial attention, shared the early temporal profile of dimension-general spatial attention but peaked and subsided earlier, suggesting that feature-specific attention precedes a more general spatial signal that drives the response.

To assess the behavioral relevance of task and distractor representations, we used multi-level models to predict single-trial RT from RSA coding strengths at each timepoint (see **Methods**). All three representations predicted faster responses, with effects arising at the stimulus onset and peaking around 450 ms (Fig. 4E, solid lines), indicating that stronger expression of task-relevant representations facilitates task performance. We also found weak but consistently positive interaction effects with parametric congruency, which suggests that the benefit of these representations was reduced when multiple distractors primed the correct response (Fig. 4E, dashed lines).

A key advantage of the MULTI is that it allows us to track decoded representations separately for each cued dimension. When split by cued dimension, robust cue, spatial attention and feature coding profiles were obtained for all dimensions (Fig. 4D). Notably, both spatial attention and feature coding were delayed for the pattern dimension, consistent with its slower visual integration, weaker behavioral interference, and later-emerging interference dynamics.

Finally, we asked whether any of these task-related representations are modulated by recent conflict history. If trial-to-trial control adaptation operates on the cued task, we should expect modulated cue or feature coding based on prior conflict. Stratifying the single-trial RSA data by the number of congruent dimensions on the previous trial revealed no evidence of modulation for either cue or feature representations (Fig. 4F). Consistent with our behavioral results, these results suggest that, rather than operating on task-relevant representations, conflict adaptation appears to be expressed elsewhere, pointing directly to the neural representations of distractors.

### Multivariate distractor representations are shaped by recent conflict

To directly examine these distractor representations, we formulated a second decoding-RSA model that jointly estimates feature-specific spatial attention for each dimension simultaneously (Fig. 5A). As before, the classifier was trained to discriminate task-relevant information only. By regressing the confusion profile onto conjunction vectors corresponding to the feature location of each non-cued dimension, as well as to the target location, we could recover distractor coding alongside target coding from the same model. In other words, this approach allowed us to partial out each dimension’s contribution from the others on every trial, despite their continuously changing locations and congruencies (Fig. 5A shows an example trial where the target is the dashed edge).

**Figure 5.**
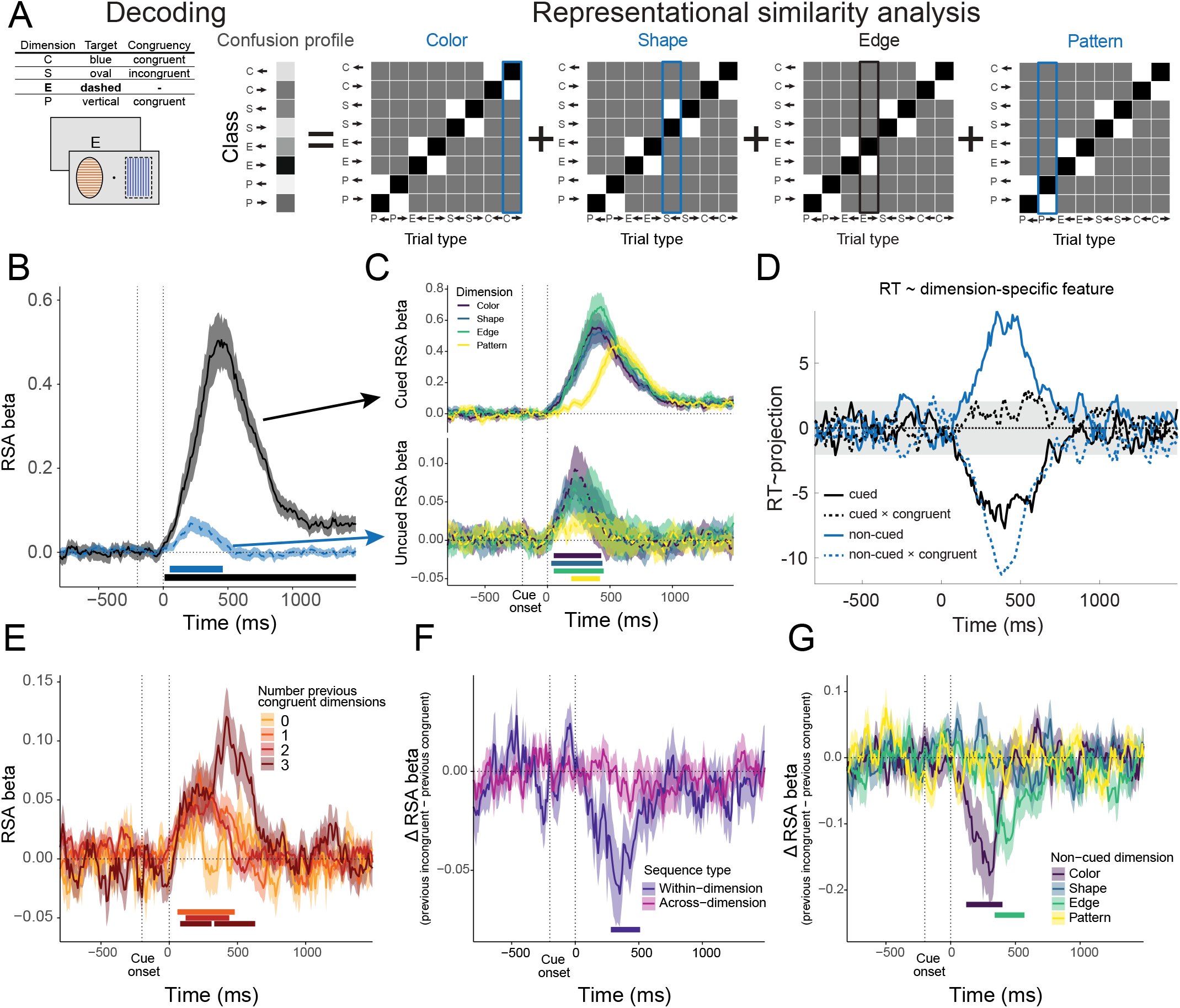
Decoding-RSA of distractor representations and their conflict-history modulation. **a**, Schematic of the distractor decoding-RSA procedure. For each trial, one conjunction vector is selected per dimension based on that dimension’s instructed feature location, yielding four trial-specific regressors that are projected against the classifier confusion profile to simultaneously estimate spatial attention directed toward each dimension’s feature. **b**, Temporal profiles of the cued (black) and average distractor (blue) representation strength. Both rise after stimulus onset, confirming parallel encoding of target and distractor features. Shaded areas indicate ±1 SEM; vertical dashed lines mark cue and stimulus onsets. **c**, Representation strength split by cued (top) and non-cued dimension (bottom). The pattern dimension exhibits notably weaker encoding, consistent with its lower behavioral interference potency. **d**, Multi-level regression coefficients show that stronger target representations predict faster RTs, while distractor representations facilitate performance when congruent and impair it when incongruent. **e**, Average distractor representations, stratified by preceding trial congruency. Distractor representations are strongest following fully congruent trials and weakest following fully incongruent trials, mirroring the behavioral CSE. **f**, Within-versus across-dimension adaptation effects on distractor representations, computed as the difference in representational strength following incongruent versus congruent trials. Robust within-dimension effects (blue) emerge around 350 ms, with no significant across-dimension effects, directly paralleling behavioral adaptation. **g**, Within-dimension adaptation effects split by distractor dimension. Effects peak earlier for color and later for edge, shape shows weaker and more diffuse adaptation, pattern shows no reliable effect.

This approach revealed robust encoding of all four dimensions in parallel (Fig. 5B). Both target and distractor (blue) representations rose after stimulus onset, confirming that non-cued features can be detected alongside task-relevant information. The expression of distractor representations was robust even when controlling for their congruency level across trials. Distinct representations were reliably detected for each distractor dimension (Fig. 5C, bottom) and each cued dimension (Fig. 5C, top), with strength scaling with behavioral interference potency.

Distractor coding peaked around 250 ms after stimulus onset, consistent with early attentional capture. These early and short-lived distractor profiles suggest a fast-acting inhibitory mechanism that suppresses distractor processing.

To confirm the behavioral relevance of these distractor representations, we regressed trial-by-trial RT on the strength of target and distractor coding using multi-level models (Fig. 5D, **Methods**). Crucially, distractor representations facilitated performance when congruent, and impaired performance when incongruent. In other words, the neural signatures we recovered directly reflect the behavioral impact of each distractor on response selection.

We next asked whether distractor representations are modulated by recent conflict history. To test this, we stratified the single-trial average distractor coding by the number of congruent dimensions in the preceding trial. This revealed a clear parametric effect: distractor representations were strongest following fully congruent trials, and weakest and more short-lived following fully incongruent trials, with intermediate levels of congruency falling along this continuum (Fig. 5E). This pattern directly mirrors the behavioral parametric CSE (Fig. 2AB), confirming that conflict adaptation is driven by the strength of distractor representations.

We next tested whether these adaptations are dimension-specific. Following the logic of the behavioral analyses, we computed adaptation effects for each combination of previous and current distractor dimension as the difference in representational strength following incongruent versus congruent trials. (Note that unlike the behavioral analysis, these effects capture a simple difference in representation strength rather than a double difference in RT.)

Adaptation effects were strongly dimension-specific. We observed robust within-dimension adaptation, marked by a negative deflection peaking around 350 ms, with no significant across-dimension effects (Fig. 5F). When split by dimension, color and edge showed the most pronounced adaptation, emerging earlier for color (peak: 300 ms) and later for edge (peak: 430 ms). The shape dimensions exhibited a weaker, more diffuse pattern, while no reliable adaptation was found for pattern (Fig. 5G). Notably, within-dimension adaptation reflected both suppression following incongruent trials and enhancement following congruent ones (Fig. S2).

Together, these results establish that distractor features are processed alongside task-relevant information, but their influence on the response selection is curtailed by a rapid, stimulus-triggered suppression. This reactive suppression mechanism operates after distractor processing has begun. However, the efficiency of reactive suppression increases with prior exposure to similar interference, effectively reflecting a form of proactive control. This pattern reveals a fundamental property of control: proactive adaptation does not prevent distractors from being encoded but instead tunes the speed and strength of the reactive suppression that resolves their interference.

### Proactive tuning of distractor suppression explains task-specific conflict learning

So far, our analyses have focused on short-term control adjustments in response to recent conflict, using only trials from *neutral* blocks, in which each non-cued dimension was equally likely to appear as congruent or incongruent. However, cognitive control can also adapt to more stable environmental regularities, allowing learned control settings to be rapidly reinstated when appropriate. To examine this long-term form of control learning, we turned to the TSPC manipulation, in which one task dimension was stably associated with a high probability (75%; MI) of distractor interference across blocks, another with a low probability (25%; MC), and the other two with a neutral probability (50%). These assignments occasionally rotated across blocks. This design established stable, learnable relationships between each task and the expected level of interference, allowing participants to anticipate control demands with sequences of task repetitions, and to adjust attention proactively.

Participants adapted control to the TSPC manipulation (Fig. 2GH). Generalized Bayesian linear regression models provided decisive evidence for a current congruency × TSPC interaction on both RT and accuracy (ERs = ∞). MI tasks showed attenuated congruency effects, reflecting reduced sensitivity to non-cued dimensions, while MC tasks showed enhanced effects, reflecting stronger distractor influence. Specifically, trials in MC tasks yielded slower and less accurate responses when current congruency was low (MC_0_ – MI_0_ = 41 ms, -4.5%) and faster, more accurate responses when current congruency was high (MC_3_ – MI_3_ = -26 ms, 0.6%).

The strength of this long-term adaptation mirrored the heterogeneous potency of interference across dimensions (Fig. 2IJ). Reliable TSPC effects were observed for color (RT: BF = 3.3 × 10^3^, accuracy: BF = 5.2), shape (RT: BF = 2.3 × 10^5^, accuracy: BF = 469), and edge (RT: BF = 57, accuracy: BF = 1.2), but not for pattern (RT: BF = 0.1, accuracy: BF = 0.1).

To determine whether TSPC effects reflect proactive or reactive implementation of control, we restricted our analyses to switch trials, the first trial of a new task sequence. On these trials, participants cannot anticipate which dimension will be cued and therefore cannot proactively load dimension-specific control settings during the ITI. If TSPC effects depend on proactive implementation, they should be absent in switch trials. If they are implemented reactively, they should emerge upon stimulus onset regardless. We found strong evidence for a PC × congruency interaction on RT (ER = 420), indicating that congruency effects were reliably attenuated for MI relative to MC dimensions already on the first trial of a new sequence. Evidence for the same interaction on accuracy was present but weaker (ER = 5.8). These findings rule out proactive retrieval of dimension-specific control settings during the ITI, pointing instead to reactive implementation upon stimulus onset.

To test whether long-term control adaptation also shaped neural distractor representations, we applied our decoding-RSA model to estimate the strength of cued and non-cued feature representations as a function of TSPC condition (Fig. 6), using a bootstrapping procedure to isolate genuine adaptation effects from trial-level congruency differences across blocks (see **Methods**).

**Figure 6.**
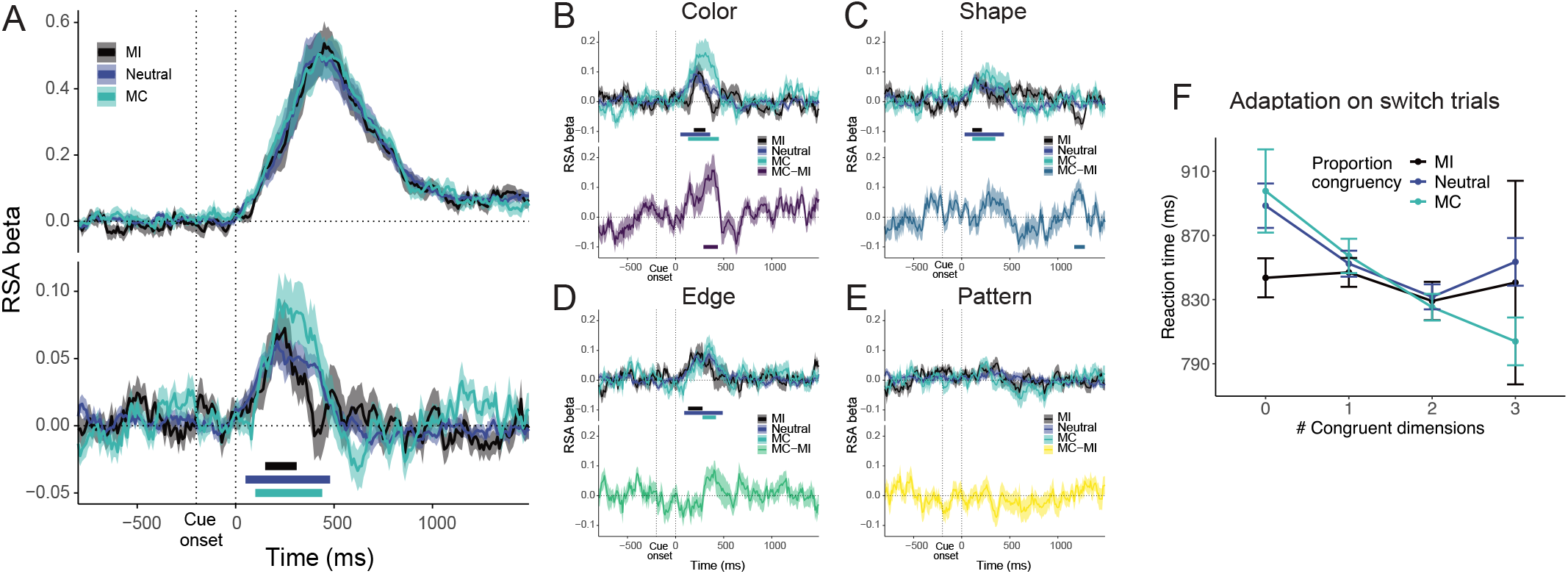
Long-term control adaptation modulates distractor representations reactively. **a**, Cued dimension (top) and distractor (bottom) representation strengths as a function of TSPC condition (MI, Neutral, MC). TSPC did not modulate target representations, while distractor representations were strongest and most sustained for MC tasks, followed by neutral and MI tasks, consistent with long-term learned modulation of distractor processing. Shaded areas indicate ±1 SEM. Vertical dashed lines mark cue and stimulus onsets. **b-e**, MC versus MI contrasts for distractor representations split by non-cued dimension: color (b), shape (c), edge (d), and pattern (e). Effects are largest for color, weaker for shape and edge, and absent for pattern, paralleling the behavioral TSPC results. **f.** TSPC effects on RT for switch trials only, on which proactive loading of dimension-specific control settings is not possible. Reliable TSPC effects nonetheless emerge, confirming reactive enactment of long-term control adaptation.

The neural dynamics of TSPC adaptation closely mirrored those of the short-term CSE. TSPC did not modulate cued dimension representations (Fig. 6A, top), confirming that long-term adaptation, like short-term adaptation, does not operate on target coding. Instead, modulation emerged exclusively in non-cued feature representations (Fig. 6A, bottom), with distractor representations strongest and most sustained for MC tasks, followed by neutral and MI tasks, consistent with reactive distractor suppression. The MC–MI contrast was largest for color, with trending effects for shape and edge, and no adaptation for pattern, paralleling the behavioral results (Fig. 6B–E).

These results demonstrate that people learn stable associations between task dimensions and control settings, allowing them to regulate distractor processing across contexts. Critically, the switch trial analysis and the temporal profile of neural distractor representations converge on the same conclusion: even control settings acquired over an extended timescale are enacted reactively upon stimulus onset. Together with the CSE results, this supports a unified account in which proactive adaptation shapes the speed and strength of a stimulus-triggered suppression mechanism.

### Conflict is encoded in dimension-specific orthogonal subspaces

The decoding-RSA approach established that distractor suppression operates through feature-based attention mechanisms and adapts to conflict history. However, it does not address a fundamental question: how is conflict itself encoded in neural activity across multiple simultaneously competing dimensions? To answer this, we applied a multivariate encoding model that directly estimates neural coding axes for conflict and spatial attention, without the constraints of the RSA approach (Fig. 7; see **Methods**). For each domain, axes were orthogonalized to separate signals shared across all dimensions from those private to each^34^, yielding four dimension-specific axes and one dimension-general axis. Distractor spatial attention was estimated by projecting these axes onto non-cued dimensions, allowing us to read out distractor-directed attention from a model trained on target-relevant information.

**Figure 7.**
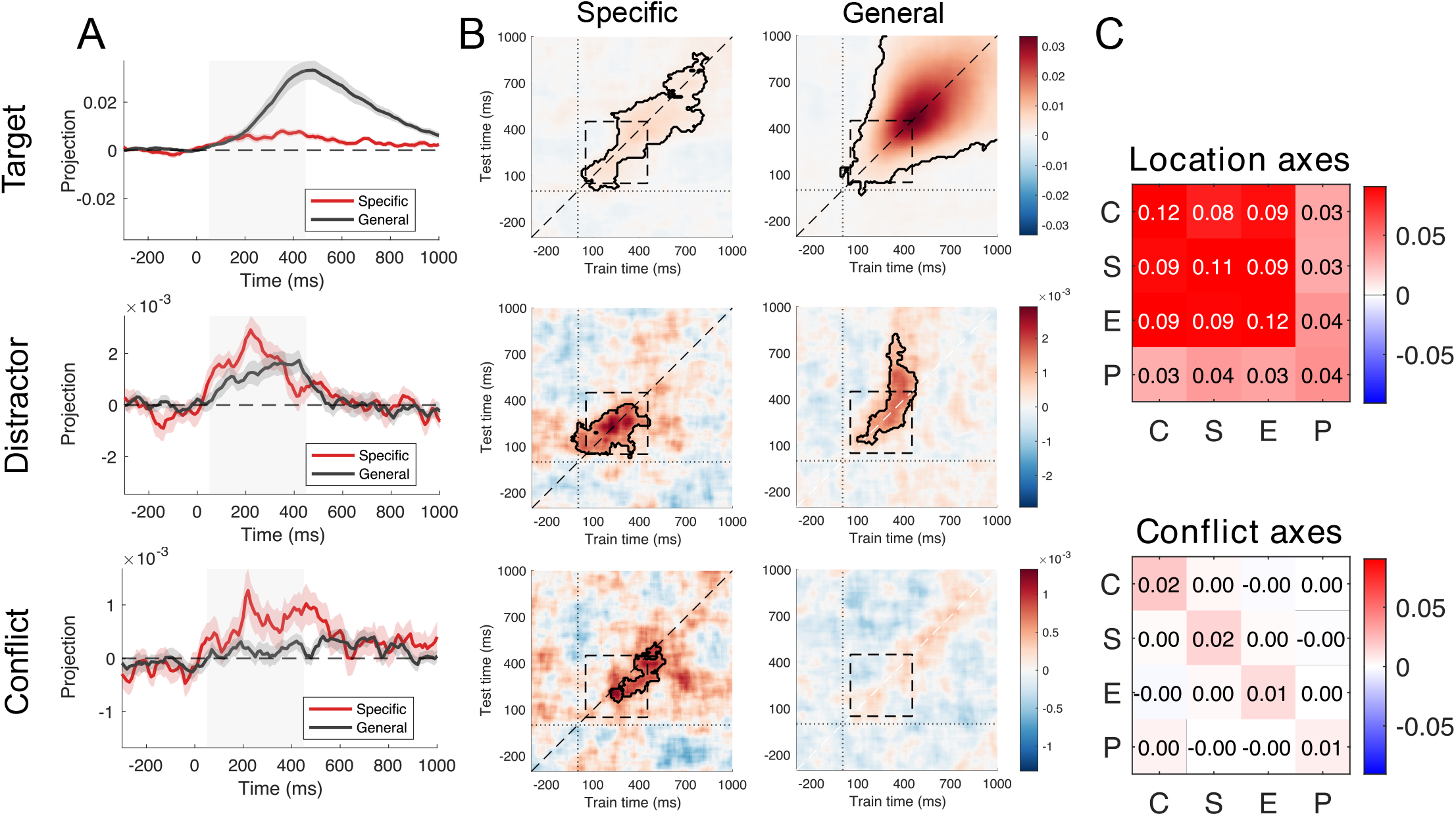
Encoding model reveals orthogonal conflict subspaces. **a**, Projections onto dimension-general (grey) and dimension-specific (red) axes for attention towards target (top), towards distractors (middle), and conflict (bottom), averaged across dimensions (mean ± SEM). Data are locked to stimuli onset (*t*=0); shaded region marks the 50–450 ms pre-response window (see Fig. 4D top). Location axes were fit using the cued dimension’s target location; distractor attention was estimated by projecting these axes onto non-cued dimensions and sign-correcting by each non-cued dimension’s feature location, allowing read-out of distractor-directed attention from a model trained exclusively on target-relevant information. **b.** Temporal generalization (TG) of dimension-specific (left) and dimension-general spatial attention (right) for the target feature (top), distractor spatial attention (middle) and conflict (bottom). Color indicates projection strength; black contours denote significant clusters (cluster-based permutation test, *p* < .01; 5,000 permutations). Dashed square marks the 50–450 ms pre-response window. **c**. Cross-dimension similarity of encoding axes, computed on the original axes prior to orthogonalization. Pairwise covariance-weighted cosine similarity across dimension axes for location (top) and conflict (bottom), averaged across the 50– 450 ms window (shaded region in **a, b**). Location axes are strongly shared across dimensions; conflict axes show near-zero pairwise similarity, indicating that each dimension occupies an orthogonal representational subspace for conflict coding.

The spatial attention axes revealed two key findings. Dimension-specific attention toward non-cued dimensions stabilized rapidly (∼250 ms after stimulus onset; Fig. 7A), indicating early encoding of distractor features in parallel with target features within dimension-specific subspaces. Second, the dimension-general component, capturing left–right spatial attention shared across dimensions, showed a striking asymmetry between target and distractor processing. When oriented to target locations (Fig. 7AB, top), this signal converged into a stable, sustained signal around response, but when oriented to distractors, spatial attention was weaker and short-lived (Fig. 7AB, middle), as if rapidly curtailed after an initial orienting response. Adaptation effects on distractor-specific and dimension-general representations largely replicated the decoding-RSA results (Fig. S3, cf. Fig. 5), while further revealing that adaptation targets primarily the shared component (spatial orienting), though dimension-specific processes are also affected for fast-integrating, high-potency dimensions such as color. In Fig. S3, response-locked analyses further confirmed that the dimension-general component peaked at response when oriented to the target, and disappearing entirely when directed to distractors, consistent with suppression preventing the stabilization of distractor-related attentional signals.

For attention axes, the shared component was dominant, as confirmed by their similarity structure (Fig. 7C). In contrast, the four conflict axes revealed near-zero pairwise cosine similarity (Fig. 7C, bottom), indicating that each dimension occupied an independent representational subspace for conflict coding. This directly challenges scalar models of cognitive control, which assume a single shared conflict signal drives control adjustments across all sources of interference.

Conflict projections were largely confined to dimension-specific subspaces (Fig. 7A, bottom), with the dimension-specific axis rising earlier and more sharply than the dimension-general axis, which was substantially weaker and more temporally diffuse. Temporal generalization analysis refined this dissociation (Fig. 7B bottom): dimension-specific conflict axes showed a temporally focused, largely diagonal pattern indicating transient, rapidly evolving coding after stimulus onset. The dimension-general axis exhibited a weak but sustained pattern, emerging around the peak of dimension-specific conflict and persisting well beyond the mean RT (∼660 ms), suggesting the shared signal outlasts the response itself. Although this pattern did not reach significance in cluster-based permutation testing, its temporal profile is consistent with a late-emerging convergence of dimension-specific conflict signals onto a shared monitoring channel. The representational geometry underlying this dissociation (Fig. 7C) suggests that conflict coding is multiplexed across dimensions, with independent monitoring signals unfolding in orthogonal subspaces before converging onto a common dimension-general representation.

## Discussion

Using behavioral, computational, and neural analyses, we show that multidimensional attention control operates through selective, reactive suppression of distractor representations. This adaptation is guided by a multivariate conflict signal encoded in orthogonal, dimension-specific neural subspaces. These findings challenge classic accounts of cognitive control that assume univariate conflict signals, and offer a new framework for understanding how the brain manages interference in complex, multidimensional environments.

Classical conflict monitoring models assume that the brain shapes attention policies using a global control signal calculated from response-or stimulus-level conflict^4,12^. Our findings challenge this account: within a single task, conflict is first represented in dimension-specific subspaces, with control tuned to individual sources of interference, and only later within the trial do these signals converge onto a shared, domain-general representation. Consistent with prior reports of both task-specific and domain-general conflict signals in medial frontal cortex^35^, and partially overlapping conflict representations across tasks in EEG^36^, we show that structured, multivariate conflict geometry operates even within a single multidimensional environment, where multiple sources of interference compete simultaneously. This pattern of adaptation suggests that the brain tracks the multidimensional statistical structure of the environment, efficiently learning control settings tailored to each source of interference, rather than adjusting a single global gain^37^.

A key insight is that attentional control operates on emerging stimulus representations rather than preventing distractors from being encoded^25,38,39^. Distractor features are processed in parallel with targets, peaking around 220 ms, precisely matching the profile of neural responses to salient distractors in visual search paradigms^23^. Distractor representations are then rapidly curtailed, with suppression strength scaling with prior conflict history, though these adjustments primarily reflected spatial attention, with feature-level adaptation reserved for high-potency dimensions, mirroring the heterogeneity in the behavioral interference effects. This timing is consistent with biased-competition frameworks^40,41^, in which top-down signals resolve competition after perceptual representations develop. Our results show that distractor features are nevertheless encoded in parallel until triggering spatial orienting, indicating that such competition and subsequent modulation can operate across distinct, spatially separated feature dimensions.

Our findings raise important questions regarding how conflict monitoring is implemented. If each source of conflict is encoded in an independent monitor, putatively by dorsal anterior cingulate cortex (dACC), how could this implementation possibly scale to handle the vast diversity of tasks and conflict sources we encounter? One possibility is that monitors are constructed ad hoc by flexibly routing strongly encoded features of the task set (e.g., cued targets, attended distractors) and performance-related variables (such as online effort estimates) into dACC, which could non-linearly expand them into mutually independent conflict subspaces^35,42–44^. Such ad hoc routing implies that inputs into dACC are dynamically reconfigured as task demands and feature salience shifts^42,43,45–48^. Another possibility is that a dedicated set of conflict monitors is embedded in the structure of dACC connectivity, forming an evolutionarily tailored basis of composable monitoring units^49–51^. These are open questions for future work.

A unifying finding across both trial-to-trial and task-level adaptation is that proactive learning does not establish anticipatory representational states, instead tuning a stimulus-triggered suppression mechanism on the fly. Our TSPC findings suggest that attentional control sets can be tied to abstract task representations, much like they can be anchored to temporal contexts or specific task features^7,52,53^. Critically, these learned associations were expressed reactively upon stimulus onset, as confirmed by the persistence of TSPC effects on switch trials, where proactive loading of dimension-specific settings is impossible. This suggests that proactive and reactive control rely on similar mechanisms, with experience setting the operating parameters under which the suppression process unfolds^54^. However, it remains possible that proactive adjustments occur at meso-or micro-levels within smaller neural populations that are difficult to detect with EEG. Future work should examine whether more reliable distractor-dimension associations can elicit sustained anticipatory neural modulation.

The control architecture uncovered by the encoding analyses may allow the system to simultaneously track multiple sources of interference without dimensional crosstalk. Dimension-specific signals provide the granularity needed for targeted distractor suppression, whereas dimension-general conflict aggregates these into a scalar readout modulating overall response caution^55^, consistent with frontal midline theta scaling with overall conflict level and peaking around the response (Fig. S1). It also remains possible that proactive adaptation manifests as phasic bursts during the ITI rather than sustained activity^56^, which would be difficult to detect with event-related EEG analyses^31–33,57,58^.

Our findings also speak to an intriguing cross-species difference. Non-human primates trained extensively on task-set conflict tasks show little interference from non-cued dimensions^34,42,59^, suggesting that highly orthogonalized feature representations emerge only after long-term consolidation. In humans, who rarely undergo such extended training60, adaptation may instead operate through modulation of representational strength within existing coding geometry, consistent with biased-competition accounts^40^. Whether geometry-based reconfiguration emerges with extended practice remains an open question. A hypothetical mechanism would be reorganizing distractor representations into subspaces more orthogonal to task-relevant dimensions^42,43,47^. Dissociating strength-and geometry-based accounts remains an open challenge, as it requires fitting encoding models directly on distractor axes, which was not attempted here for low signal-to-noise in distractor-level representations.

## Methods

### Participants

We recruited 27 participants (22 females, 5 males, 1 nonbinary; mean age: 27 years, range: 20-39) with no history of neurological disorders and normal or corrected-to-normal vision. Recruitment was restricted to right-handed participants without color blindness. Participants were recruited from the St. Louis community via the Washington University School of Medicine Research Participant Registry and from the Washington University in St. Louis student participant pool. All were compensated with $25. The experimental procedure was approved by the Institutional Review Board of Washington University in St Louis (IRB #: 202110190). Written informed consent was obtained before the experiment, and participants were debriefed afterwards. Sex or gender was not analyzed, as it was not pertinent to our hypotheses. We aimed for a sample size of about 25 participants based on the size of behavioral effects observed in our previous work^17^. However, no prior effect size information was available for the new EEG effects; therefore, no statistical method was employed to define the sample size. One participant was excluded from the analyses due to partial data recording.

### Behavioral procedure

To study attentional control over multiple independent stimulus dimensions, we used the MULtidimensional Task-set Interference paradigm^17^ (MULTI; Fig. 1), implemented in Psychtoolbox^61^. On each trial of the MULTI, participants attend to a letter cue and choose between two objects, presented side by side, that differ in four dimensions: color, shape, edge, and pattern. For each of these dimensions, there are two features (blue/orange for color, oval/rectangle for shape, solid/dashed for edge, and horizontal/vertical for pattern), which are randomly assigned to the left and right objects on each trial. Each object, therefore, consists of one feature from each dimension, and these features are fully randomized across trials. All features are highly discriminable, and the objects are matched for size.

The letter cue indicates which dimension is task-relevant on a given trial. Participants’ task was to identify which object contained the target feature for that dimension and select its side by pressing a corresponding button. At the beginning of the experiment, participants were instructed on which feature was the target for each of the four task dimensions. For instance, a participant might be instructed to choose the object with blue color if the cue is C, the oval shape when it is S, the dashed border when it is B, or the vertical pattern when it is P. The specific target features were randomly assigned for each participant and remained constant throughout the task.

The cued task dimension switched in pseudo-randomized sequences, making each dimension periodically relevant. As in ^17^, the same dimension was cued for 3 to 5 consecutive trials, before switching to another. All 4 dimensions were cued within 4 switch trials. Therefore, participants constantly cycled through the 4 tasks, organized in a series of 3 to 5 repeat trials. This ensured that all dimensions were cued an equal number of times, and that each of them had been cued relatively recently at each point in the task.

For suitability with EEG data collection, some changes were introduced to the task and trial structure compared to our prior work. Most importantly, the cue and stimulus displays were temporally separated, preventing overlap. Each trial started with a brief presentation of the letter cue (50 ms), followed by a fixation dot (150 ms). Next, the two objects appeared, flanking the fixation dot, and remained on screen until response or until a response deadline of 2 s had passed. Participants indicated their choice by pressing the left or right button on a Cedrus RB-840 response pad (Cedrus Corporation, San Pedro, CA, USA). An incorrect or missing response was immediately followed by negative feedback, presented as a red “x” at the center of the screen for 1 s. The inter-trial interval randomly varied between 1.3 and 1.5 s with a fixation cross displayed throughout.

Participants performed 32 blocks of 50 trials each (1600 trials in total), separated by self-paced breaks. Before the main task, participants performed five blocks of practice. These included four blocks in which participants learned to respond to each dimension individually, and a fifth block in which they practiced switching between task dimensions. These practice blocks ensured that participants had mastered the cue–feature mappings before EEG recording began.

In line with the previous design, task-set interference was parametrically manipulated on a discrete congruency scale from 0 to 3. On each trial, any of the three non-cued dimensions could be congruent if they primed the same response side as the cued target feature, or incongruent if they primed the opposite side. This manipulation enabled us to examine trial-by-trial adjustments in attentional control as a function of recent multidimensional conflict.

To investigate sustained associations of control settings with task dimensions, we introduced a task-specific proportion congruency (TSPC) manipulation^7^. This manipulation established stable relationships between cued task dimensions and expected conflict levels. At each point in the experiment, one task was mostly incongruent (MI; i.e., each of the three distractors had a 25% probability of being congruent), one task was mostly congruent (MC; 75% probability), and the remaining two were neutral (50% probability). These assignments rotated three times across the session according to a Latin Square design, with each TSPC mapping lasting eight blocks of 50 trials. This rotation ensured that, across the experiment, each task dimension served once as MI and once as MC, providing a fully counterbalanced design. For each participant, the initial assignment of PC to each task dimension was randomized.

Participants were instructed to respond as quickly and as accurately as possible. To minimize eye and head movements, they were requested to always maintain fixation at the center of the screen and to process the left and right objects using peripheral vision. The average duration of the experiment was 2.5 hours, including approximately 1 hour for EEG setup and instructions, 1 hour for task execution, and the remaining time allocated for debriefing and the cleaning of the EEG cap and electrodes.

### Behavioral data analysis

Our behavioral analyses followed the procedure reported in our previous behavioral investigation using the MULTI ^17^. Here, we provide a synthesis of that procedure. Switch trials were excluded from the analyses. For congruency sequence effect (CSE) analyses, we also excluded trials following incorrect responses to avoid contamination from post-error slowing^62^ or other control processes. For proportion congruency analyses, all trials were included. Incorrect trials were further excluded for all RT analyses.

Accuracy and reaction times were modelled using a Bayesian multilevel generalized linear approach implemented in R ^63^ with the ‘brms’ package^64^. Models predicting reaction times assumed a shifted lognormal response distribution, with identity and log link functions for the distributional parameters. Models predicting accuracy assumed a Bernoulli distribution with a logit link function. We report medians and 95% credible intervals of the posterior parameter distributions on the log scale for reaction times, and the logit scale for accuracy, and estimated marginal means are reported in the response scales (milliseconds for reaction times and percentages for accuracy). All models used weakly informative priors centered on zero for the parameters of interest and on the raw mean for the intercept. For hypothesis testing, we report evidence ratios (ER).

To assess within-and across-dimension adaptation, we calculated CSEs for 16 possible sequences of non-cued dimensions. Each sequence compared the change in interference for a given dimension after congruent versus incongruent trials on another dimension. Some sequences assess within-dimension adaptation (e.g., previous color congruency on current color sensitivity), while others assess across-dimension adaptation (e.g., previous shape congruency on current edge sensitivity). For each combination, we computed the adaptation effect by subtracting the interference effect after incongruent trials from the interference effect after congruent trials.

For the TSPC analyses, we modeled RTs and accuracy as a function of current congruency, TSPC condition, and their interaction. This allowed us to test whether the magnitude of the congruency effect varied as a function of each task’s current congruency bias (MI, MC, neutral).

### Neural network modeling of conflict adaptation

To capture the temporal dynamics of dimension-specific control adaptation, we implemented a dynamic neural network model of the MULTI (Fig. 3A), inspired by classic architectures of control allocation and conflict monitoring in interference paradigms^4,65^ and building on the static model introduced by Gheza and Kool^17^. The model simulates how evidence from multiple dimensions accumulates over time and how recent conflict shapes pathway-specific control.

The network consists of four parallel pathways, corresponding to the four dimensions of the task. Each pathway begins with two input units encoding the side of the target feature for that dimension (left vs. right). Each set of input nodes then projects to two units in the hidden layer. These units integrate input from both the stimulus units and a set of task units representing top-down attention towards each dimension. The task units modulate the gain of their corresponding hidden units by offsetting a fixed negative bias, thereby enhancing sensitivity to stimulus input in the attended dimensions. The activation of each task unit depended on whether its corresponding dimension was cued and was further adjusted based on recent dimension-specific conflict (see below). This attention gating mechanism increased the separation between the activations of hidden units for attended dimensions, as the activity of the task units offset the inhibitory bias, allowing hidden units to encode stimulus input with more sensitivity. The units in the hidden layer project to two response nodes (left vs. right) via balanced excitatory and inhibitory connections. Specifically, hidden units representing target features on the left projected positively to the left response node and negatively to the right response node, and vice versa.

Net input into every hidden and response unit was computed as the sum of the activity of projecting units multiplied by their corresponding connection weight, with an added constant bias. This resulting net input was then passed through a logistic activation function, which converted it into an activity value between 0 and 1.

To simulate temporal dynamics, unit activations at the hidden and response layer evolved over time using leaky integration^66^. At each timestep, the activity of the hidden layer was a weighted combination of its previous activation and its current net input, controlled by a pathway-specific time constant (*τ*_*d*_). For each timestep *t*, the net input *H* into each hidden neuron *n* in dimension pathway *d* accumulates a small dimension-specific trickle of evidence:

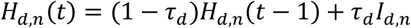

with *H*_*d,n*_(0) = 0 ∀ *d, n*, and where *I*_*d,n*_ is the total net input to neuron *n* in dimension pathway *d*. This input is constant within a trial and is the sum of stimulus input *S*, task input *T*, and bias term *b*_*H*_:

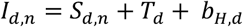

This produces dimension-specific temporal integration without lateral interactions at the hidden layer. To simulate the choice process, the two response units likewise integrated their inputs using a leaky integrator:

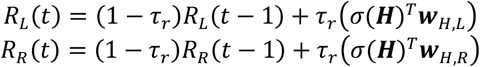

with *R*_*L*_ and *R*_*R*_ the activity of the left and right response units, *τ*_*r*_ a response-level time constant, ***H*** the activity of the hidden layer at timestep *t*, ***w***_*H,L*_ and ***w***_*H,R*_ the connections between the hidden layer and the right and left response unit, and *σ* the logistic activation function.

The model’s choice mechanism consisted of two opposing evidence accumulators. At each timestep, the instantaneous evidence was proportional to the difference between the two response activations Δ(*t*), which drives a pair of opposing noisy accumulators:

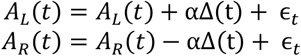

with Gaussian noise ϵ_*t*_∼*N*(0, σ). The trial terminated when either accumulator reached a fixed decision threshold *θ*. The first unit to cross this threshold determined the model’s choice, and the corresponding timestep defined the reaction time for that trial. Accuracy was defined by the match between the selected response and the cued target feature’s location.

To simulate conflict monitoring, we followed and extended our prior modeling work. As before, we computed distractor-specific conflict *E*_*d*_ as Hopfield energy ^26^, but now for each time point *t*:

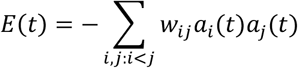

where *w*_*ij*_ represents the connection weight between units *i* and *j, a* indicates unit activity and both subscripts are indexed over all units in the pathway. Here, we compute this measure of energy between the hidden and response layer separately for each non-cued dimension.

Because of the excitatory and inhibitory configuration of the connections between hidden and response units, the sign and magnitude of the dimension-specific conflict are only a function of the pathway state. If dimension-specific conflict is positive, a non-cued dimension provides evidence inconsistent with the correct response, and negative if it provides consistent evidence

Next, for each dimension, conflict is transformed into a measure of control using the following formula:

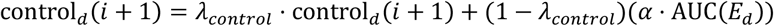

where

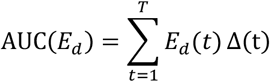

indicates discrete integration of pathway-specific energy across the trial, and with *i* indexing trials, *λ*_*control*_ allowing a time-discounted measure of previous conflict, and *α* a scaling parameter. At the start of each trial, each non-cued dimension’s control value was then subtracted from the (zero) input of that dimension. This algorithm implements distractor-specific adaptation, with the model becoming more sensitive to a non-cued dimension after a congruent trial on that dimension, and less sensitive after an incongruent trial.

Model parameters (including connection weights) were hand-tuned within plausible ranges to reproduce the key qualitative behavioral patterns (interference, dimension-specific adaptation, etc.). Similar qualitative results were obtained across a broad range of parameter values, confirming that the reported dynamics do not depend on specific fine-tuned settings.

### EEG recordings and preprocessing

EEG data were recorded using a 64-channel Brain Vision actiCHamp Plus system (Brain Vision LLC), placed on a lycra cap (actiCap) following the 10/20 system, and sampled at 500 Hz. Data were collected from 61 scalp sites (Fpz, Fp1, Fz, F3, F7, FC1, FC5, Cz, C3, CP1, CP5, Pz, P3, P9, PO3, PO7, Oz, PO4, PO8, P4, P10, CP2, CP6, C4, FC2, FC6, F4, F8, Fp2, AFz, AF3, AF7, F1, F5, FCz, FC3, FT7, C1, C5, T7, CP3, P1, P5, P7, O1, POz, O2, P2, P6, P8, CPz, CP4, C2, C6, T8, FC4, FT8, F2, F6, AF4, AF8). The Cz electrode served as the online reference site. Three electrodes were placed lateral to the external canthi and below the right eye for measuring horizontal and vertical eye movements (electro-oculogram, EOG), and the ground electrode was placed at channel location AFp1.

The preprocessing pipeline was developed with EEGLab v2021.1 ^67^. It contained the following steps. The continuous signal was filtered offline with separate high-pass (0.1 Hz lower edge frequency) and low-pass (80 Hz higher edge frequency) Hamming windowed sinc FIR filters. Independent Component Analysis (ICA) was used to identify eye artifacts generated by blinks and saccades, muscle activity, and channel noise. To maximize component separation ^68^, a copy of the data was high-pass filtered at 2.0 Hz and segmented into epochs time-locked to objects onset (-500 to 1500ms) for ICA training; the resulting ICA weights were then transferred to the original dataset. Artefactual components were manually identified by inspecting activation time courses, topography, power spectrum, and with the aid of an automated classifier ^69^.

Next, stimulus-locked epochs were extracted in a larger interval (-950 to 2050 ms) suited for time/frequency analysis, and the non-artefactual independent components were back-projected to electrode space. Baseline activity (mean voltage during the -500 to -300 ms window) was subtracted from each epoch. An automatic artifact rejection procedure was applied to discard trials with i) voltage exceeding ± 120 µV in the -500 to 1500 ms interval or ii) improbable data based on joint probability of electrode activities.

Clean data were re-referenced to the average channel activity, and the signal at Cz was reconstructed. Trials with incorrect or missing responses were excluded from further EEG analyses. For congruency sequence effects, we further excluded switch trials and trials preceded by incorrect responses.

### Time/frequency analysis

Time/frequency analysis of the single-trial EEG data was performed via Morlet wavelet convolution^70^. We convolved the EEG signal with a set of complex Morlet wavelets, defined as complex sine waves tapered by a Gaussian window 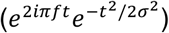, where *i* is the imaginary operator, *t* is time in seconds, *f* is frequency in Hz, and *σ* is the width of the Gaussian window. The width of the Gaussian was defined as 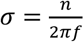, where *n* is the number of sine cycles. Because *n* governs the trade-off between temporal and spectral precision, it was set to increase logarithmically from 2 to 10 with increasing wavelet frequency. The frequencies of the wavelets ranged from 1.5 to 40 Hz in 20 logarithmically spaced steps. It follows that the full-width at half-maximum (FWHM) ranged from 500 to 94 ms with increasing wavelet peak frequency^71^.

The convolution was performed in the frequency domain and converted to the time domain using an inverse Fourier transform. Instantaneous power estimates were obtained by taking the squared magnitude of the resulting complex-valued signal, and decibel-converted by normalizing with a channel-and frequency-specific baseline. This baseline was defined as the average power across all trials and across 100ms of the inter-trial interval (-800 to -700 ms pre-stimulus, i.e., -600 to -500 ms before the cue onset). The power time series were down-sampled to 100 Hz to reduce the computational load for subsequent analyses.

### Decoding-RSA

To link EEG patterns to temporally evolving task representations, we used a representational decoding framework that captures how neural activity encodes information about attended and unattended stimulus dimensions. This approach allowed us to quantify the emergence and temporal dynamics of both target and distractor representations within each trial.

For each trial and time-point, we decoded target-and distractor-relevant information using a two-step procedure consisting of 1) time-resolved multivariate pattern classification (‘EEG decoding’), and 2) representation similarity analysis based on the resulting classification profiles (‘RSA’). This method (‘decoding-RSA’) was adapted from Kikumoto and Mayr^20^ for use with the MULTI, incorporating several modifications to accommodate its multidimensional structure and frequency-resolved EEG data.

#### Decoding

As a first step, we trained linear decoders to classify each trial based on the eight possible conjunctions of task cue (e.g., the letter ‘C’ for color) and the side of the corresponding instructed feature (e.g., left). These conjunctions define the target feature that participants search for on each trial (e.g., the color blue). The classifier thus aimed to decode which of the possible eight cue × side conjunctions was present in each trial from EEG activity at each time point.

Specifically, we trained independent classifiers for every time point using a Penalized Linear Discriminant Analysis using MATLAB’s ‘fitcdiscr’ function (*Version: 9*.*12*.*0, R2022a)*. The classification feature space was obtained by concatenating the instantaneous EEG power across all frequency bands and the 61 electrodes. To increase the signal-to-noise ratio, the original 20 frequencies were averaged within the bands Delta (1.5-3.5), Theta (3.6-7.5), Alpha (8-12.5), Beta (13-30), and Gamma (31-40), leading to 305 channel-band features. This approach generates eight-element vectors of probabilities reflecting the likelihood of a given trial belonging to each of the classes at each time point within the epoch.

We adopted a multivariate classification approach, training classifiers to separate the eight classes simultaneously, rather than multiple pairwise classifiers. This strategy was essential because it provides probabilistic outputs for all classes. We used these class probability vectors in the subsequent RSA analyses.

We evaluated classifier performance with a repeated *k*-fold cross-validation procedure. For each participant, single-trial EEG data were randomly partitioned into six “folds” with an equal number of observations for every combination of the eight classes and four PC conditions. To ensure balanced training sets, trials exceeding the minimum number of observations across these thirty-two combinations were randomly discarded. At each step of the cross-validation, the classifier was trained on five folds and tested on the remaining fold and discarded trials. This was repeated until all folds served as the test set. This cycle was repeated eight times. Critically, the trial order was shuffled before each cycle, so that each set of folds included a different, random split of the data. This repetition increased the reliability of decoding estimates and maximized trial usage despite the heavy subsampling inherent in the k-fold balancing process (resulting in approximately 50% of trials used in the training per iteration). At each step, the EEG features in train and test trials were submitted to multivariate noise normalization (whitening) by means of a covariance matrix estimated on the full epoch^73^. During training, the LDA hyperparameter Gamma (shrinkage) was optimized once per time sample within the epoch and then fixed for all following k-fold steps.

For each time sample, a trained classifier was tested on each single trial in the test set. The classification probabilities estimated for each trial were then averaged across all folds and cycles, yielding an eight-element vector for each trial and time sample. Because these classification probability vectors are compositional (i.e., the class probabilities sum to one), a centered log-ratio transformation ^74^ was applied before the subsequent RSA regression analyses.

#### RSA

In the second step, we used RSA to quantify how the probability profiles produced by the decoders reflect different task representations. For each trial and time point, the classification probability vector was simultaneously regressed onto multiple model vectors, each representing a hypothesized component of task structure. The resulting regression coefficients indexed the strength of neural representations for each component at that moment in time.

We first defined an RSA model (m1) to estimate representations of cued dimensions (Fig. 5). This model included three regressors corresponding to 1) the task “cue”, 2) the “side” of the instructed feature, and 3) the “cue x side” conjunction (the dot product of cue and side vectors), representing the conjunction that defined the target. These model vectors allowed the estimation of the neural coding of the active task rule, the location of spatial attention, and the spatial location of the target feature for the cued dimension on each trial; because target features are fixed across the experiment, this conjunction captures feature-selective spatial attention rather than feature identity per se, and is labeled ‘Feature coding’ to reflect the feature-specific nature of this spatial selection signal. Note also that, because we only analyzed correct trials, the “side” vector equally represents the location of spatial attention and response choice. Fig. 2A illustrates the RSA procedure for m1 for a given trial, depicting how the model vectors included as RSA regressors depended on the trial type (x-axis).

The trial type is defined by the location of the instructed feature for the cued dimension (i.e., the target feature). To ensure comparable scaling across regressors, the three model vectors were normalized using their Frobenius norms, and the cue and side matrices were rescaled relative to the conjunction matrix.

A second RSA model (m2) was developed to estimate dimension-specific feature representations of both cued and non-cued dimensions simultaneously. To achieve this, the RSA model included four “Cue x Side” vectors (one per dimension), each representing the location and identity of the instructed feature for a given dimension. For the cued dimension, the encoded side corresponded to the correct response, whereas for each non-cued dimension it depended on the congruency of that dimension. Because congruent and incongruent trials were unevenly distributed across task conditions, subsequent analyses used bootstrapping procedures to ensure unbiased estimates of distractor dimensions (see *Bootstrapping procedure* below).

#### Brain-behavior analyses

To assess the behavioral relevance of task and distractor representations, we used multi-level models to predict single-trial RT from RSA coding strengths (Figs. 4E, 5D). For each time point within the RSA profile, we used multi-level models to predict RT from the strength of the Cue, Spatial attention, and Feature coding strengths, and their interaction with parametric congruency. For task representations, predictors included the Cue, Spatial attention, and Feature coding strengths, their interactions with parametric congruency, and nuisance predictors (task cue, response side, and their interaction) to account for systematic RT differences across conditions. For distractor representations, predictors included the RSA betas for the target feature and the average of the three distractor features, as well as their interactions with parametric congruency and the same nuisance predictors.

### Encoding models

The decoding-RSA approach is inherently limited in the number of predictors that can be simultaneously included as regressors, given the low degrees of freedom in the eight-element LDA classification profile. For example, including four Cue × Location vectors in RSA m2 allowed simultaneous estimation of cued and non-cued dimension representations, but prevented simultaneous estimation of variance explained solely by spatial attention. More broadly, this approach cannot speak to how conflict itself is encoded across multiple simultaneously competing dimensions. To address these limitations, we developed a complementary encoding approach inspired by targeted dimensionality reduction analyses in animal electrophysiology^34^, applied here to our EEG data (channel × frequency) as the feature space.

The encoding model comprised sixteen regressors fitted simultaneously at each timepoint via multivariate linear regression: four cue regressors (one per dimension, dummy-coded), four dimension-specific spatial attention regressors (coding the left/right location of each dimension’s target feature), four dimension-specific conflict regressors (contrast-coded congruency for each non-cued dimension); four previous-conflict regressors were additionally included to partial out carry-over effects from preceding trials, but did not reveal reliable information and are not discussed further. The cue regressors were mean-centered to remove intercept-like variance before orthogonalization. The model was fitted on noise-whitened EEG data using a cross-validated scheme (6-fold, repeated across 8 random shuffles; see *Decoding*), with axes estimated on training data and projected onto held-out test data.

To obtain de-mixed trajectories separating signals common to all dimensions from those private to each, a second step applied targeted orthogonalization^34^ to the spatial attention and conflict axes. For each domain, a dimension-general axis was computed by averaging across the four dimension-specific axes; this shared component was then residualized from each dimension-specific axis, yielding four mutually orthogonal dimension-specific axes and one dimension-general axis per domain — the former capturing representational spaces private to each task dimension, the latter a common signal shared across all dimensions regardless of which was cued. The full set of resulting axes (four cue, four dimension-specific attention, one dimension-general attention, four dimension-specific conflict, and one dimension-general conflict) were pre-whitened using the residuals from the initial GLM fit. Previous-conflict axes were included in the model but not analyzed further.

Critically, whereas spatial attention axes were estimated using the cued dimension’s target location to define the regressors, distractor spatial attention was recovered by projecting these same axes — both dimension-general and dimension-specific — onto non-cued dimensions, using the location of each non-cued dimension’s instructed feature as the rectifying contrast. This allowed recovery of distractor representations from a model trained exclusively on target-relevant information.

For temporal generalization analyses (Fig. 7B), axes were estimated independently at each training time point and projected onto held-out data at all test time points, yielding a training time × test time generalization matrix. For cross-condition generalization analyses (Fig. 7C), pairwise cosine similarity between all axes was computed on the original pre-orthogonalization axes, in a cross-validated manner: encoding axes were estimated independently on two folds, residuals were pooled across folds to estimate a single noise covariance matrix via Ledoit-Wolf shrinkage regularization, and covariance-weighted cosine similarity was computed between the two sets of axes.

### Bootstrapping procedures

We implemented bootstrapping resampling procedures to prevent biased estimation of Location representations, which could arise because this coding axis was shared across dimensions. Separated procedures were used for the CSE and TSPC analyses.

For the CSE analyses (using either RSA m2 or encoding models), we restricted all analyses to the neutral TSPC blocks to avoid confounding sequential adaptation with task-specific adaptation. In addition, we downsampled the single-trial results to match the number of congruent and incongruent trials for each distractor dimension in both the current and preceding trials. This ensured balanced contributions to trial-by-trial adaptation estimates.

For TSPC analyses, block-wise differences in congruence proportions (MI vs. MC) would distort model estimates if trials were simply averaged. To counter this, we applied a bootstrap procedure to the single-trial RSA results, restoring unbiased proportions of congruent and incongruent distractors within each task dimension and PC condition. Specifically, when computing condition averages, we oversampled trials from the less frequent congruency category until both categories were equally represented.

### Significance testing on EEG results

Statistical significance over RSA and encoding timeseries was assessed using cluster-based permutation tests ^75^, implemented in MATLAB. We performed 10^4^ permutations for each contrast, with cluster *α* = 0.05. For time-frequency analyses of FMT power we identified significant clusters (*α* = 0.01) using the *EEGLAB* package ^67^, after averaging over channels of interest.

## Acknowledgements

We thank Todd S. Braver, Julie M. Bugg, the members of the Control and Decision Making Lab, and the Connections with Control group for thoughtful discussions and feedback. Wouter Kool was supported by an Office of Naval Research / Department of Defense (ONR/DoD) grant (Grant N00014-23-1-2792). Davide Gheza was supported by a McDonnell Center for Systems Neuroscience grant.

