## Supplementary Information for "Reactive suppression of distractor representations resolves multidimensional interference"

### 1. Frontal midline theta indexes multidimensional conflict and control allocation

Frontal midline theta oscillations (FMT; 3–8 Hz) are a hallmark of cognitive control, reliably linked to conflict detection and control engagement under stimulus- and response-level interference[1–3]. Whether the same mechanisms are engaged under multidimensional conflict remains an open question. Analyzing FMT therefore provides a converging test for conflict monitoring and control recruitment in the MULTI.

FMT power increased following stimulus onset and peaked around the response (Fig. S1). At frontocentral electrodes, theta power was robustly modulated by current congruency (Fig. S1C, top), replicating classic signatures of conflict detection[3]. Single-trial regressions revealed a topographic dissociation: frontocentral theta predicted slower correct responses, replicating the canonical link between midfrontal theta and conflict-driven response slowing, whereas parietal and lateral-occipital theta predicted faster correct responses (Fig. S1D). We also observed marginal effects of previous conflict at occipitoparietal channels (Fig. S1C, bottom), suggesting spatially distinct contributions of theta to current conflict detection and adaptation to prior conflict.

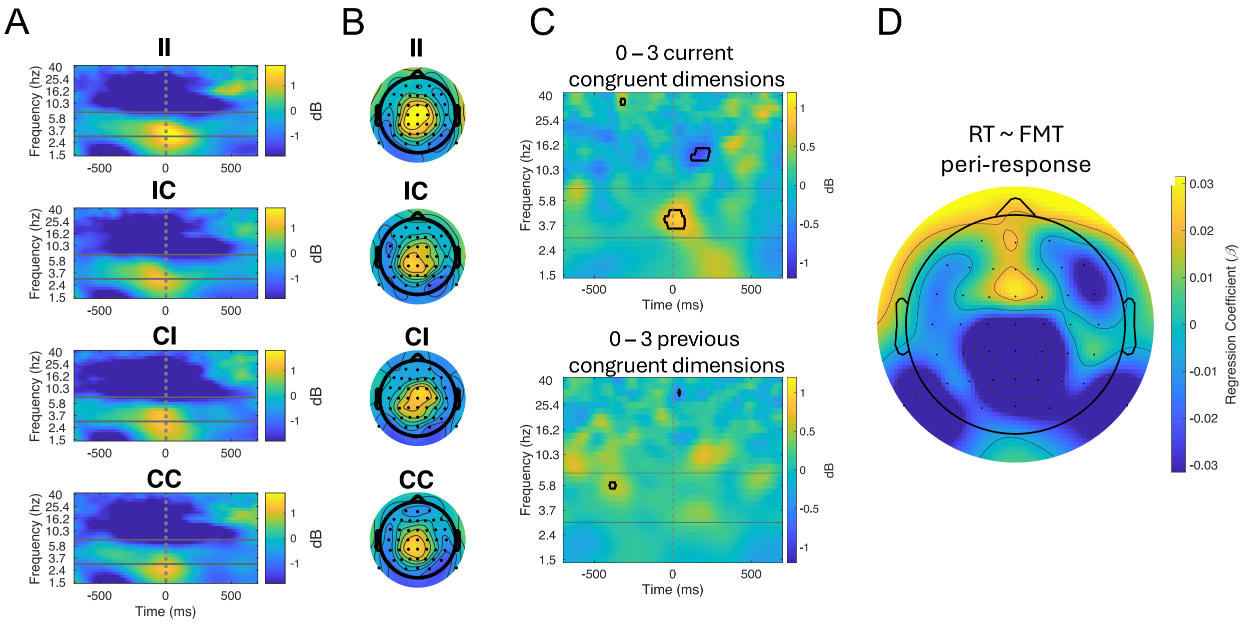


**Figure S1. a,** Time–frequency plots of response-locked power at frontocentral electrodes for four trial-sequence types (II, IC, CI, CC), where the first letter indicates congruency in the previous trial and the second letter the congruency in the current trial; "I" and "C" correspond to trials with 0 and 3 congruent dimensions, respectively. Both II and CI sequences elicited elevated theta power, with stronger activity for the former sequence type. In contrast, CC trial sequences elicited the weakest theta power. Data were averaged over frontocentral channels: Cz, FCz, FC1, FC2, C1, C2, CPz, CP1, CP2. Theta (3–8 Hz) power peaks around the response, with II sequences showing the strongest activity and CC the weakest. **b,** Corresponding scalp topographies of theta power (3–8 Hz, -75 to 75 ms peri-response window), revealing frontocentral modulation as a function of congruency sequence. **c,** Cluster-corrected time-frequency contrasts (alpha = 0.01, 5000 permutations) showing significant modulation of theta by current congruency (top) and previous congruency (bottom). The top panel shows effects averaged over frontocentral channels (Cz, FCz, FC1, FC2, C1, C2, CPz, CP1, CP2), highlighting robust theta increases for incongruent trials. The bottom panel shows effects over parieto-occipital channels (Pz, POz, CPz, CP1, CP2, P1, P3, P5, PO3, PO7, O1, P9, P2, P4, P6, PO4, PO8, O2, P10), indicating weaker but detectable influences of prior conflict. A-C. Data were response-locked and baseline-normalized using the –800 to –500 ms prestimulus window. Only trials that met all of the following criteria were included: correct responses, following a correct response, not being switch trials, and associated with neutral TSPC. **d,** Topography of regression coefficients linking peri-response (-200 to 0 ms) theta power to single-trial RTs. Higher theta at frontocentral sites predicts slower responses, whereas parietal and lateral-occipital theta predicts faster responses, consistent with conflict monitoring and control implementation processes, respectively. All correct trials were included in regression models. Single-trial power was not baseline-normalized but z-scored across trials and frequencies. RTs were also z-scored.

### 2. Conflict adaptation bidirectionally modulates distractor representations

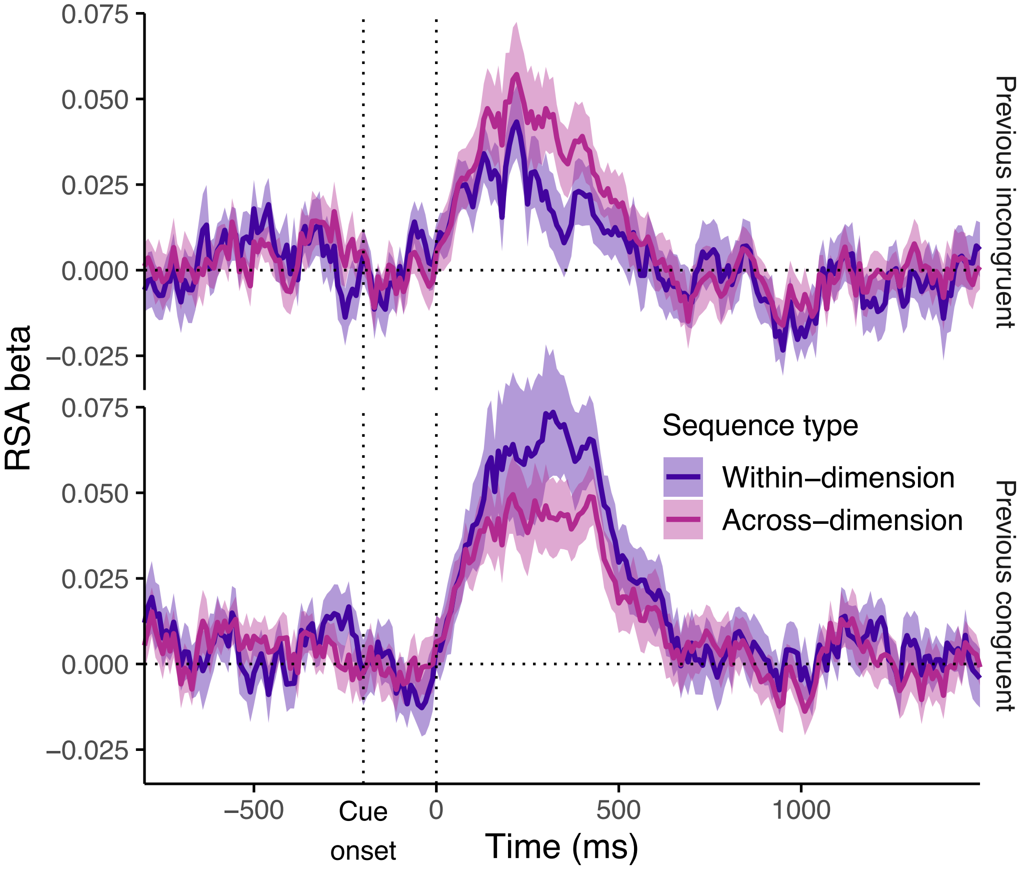


**Figure S2. Bidirectional within-dimension adaptation of distractor representations.** RSA distractor representation strength split by preceding trial congruency (incongruent, top; congruent, bottom), shown separately for within-dimension and across-dimension sequences. As can be seen in Fig. 5F, across-dimension effects are absent between previous congruency types. Following incongruent trials, within-dimension distractor representations are suppressed compared to this baseline, whereas following congruent trials, they are enhanced. This indicates that the adaptation effects in Fig. 5F reflect bidirectional modulation rather than suppression alone. Shaded areas indicate ±1 SEM; vertical dashed lines mark cue and stimulus onsets.

#### 3. Dissociable contributions of adaptation on attentional orienting and feature representations

We applied the encoding model to examine the temporal dynamics of dimension-general and dimension-specific spatial attention, the latter capturing processes private to each dimension. Even though the dimension-specific component cannot fully dissociate feature-specific spatial coding (e.g., “color-left”) from spatially invariant feature representations (e.g., “blue”), it likely captures both; we therefore refer to it as “feature coding” and to the dimension-general component as “spatial orienting” throughout.

To gain a more granular understanding of adaptation effects observed with RSA (Figs. 4 and 5), we analyzed adaptation effects on each component, spatial orienting and feature coding, separately for cued and non-cued dimensions. For cued dimensions (Fig. S3A, top), the evolution of spatial orienting resembled that obtained with decoding-RSA. In contrast, orienting towards non-cued dimensions (Fig. S3A, bottom) was markedly weaker and short-lived, consistent with rapid attentional capture followed by reactive suppression. When we examined trial-by-trial adaptations in this spatial orienting signal, we replicated a reliable within-dimension effect, with no appreciable effect across-dimension (Fig. S3BC). Moreover, we saw significant within-dimension adaptation effects for color, shape, and edge (the same dimensions that exhibited behavioral adaptation; Fig. 5G; cf. Fig. 2C-F), whereas pattern showed no measurable modulation, consistent with its later-peaking (Fig. 3A) and therefore weaker interference (Figs. 1FG). The timing of adaptation effects for the remaining dimensions tracked their faster interference dynamics (Fig. 3A).

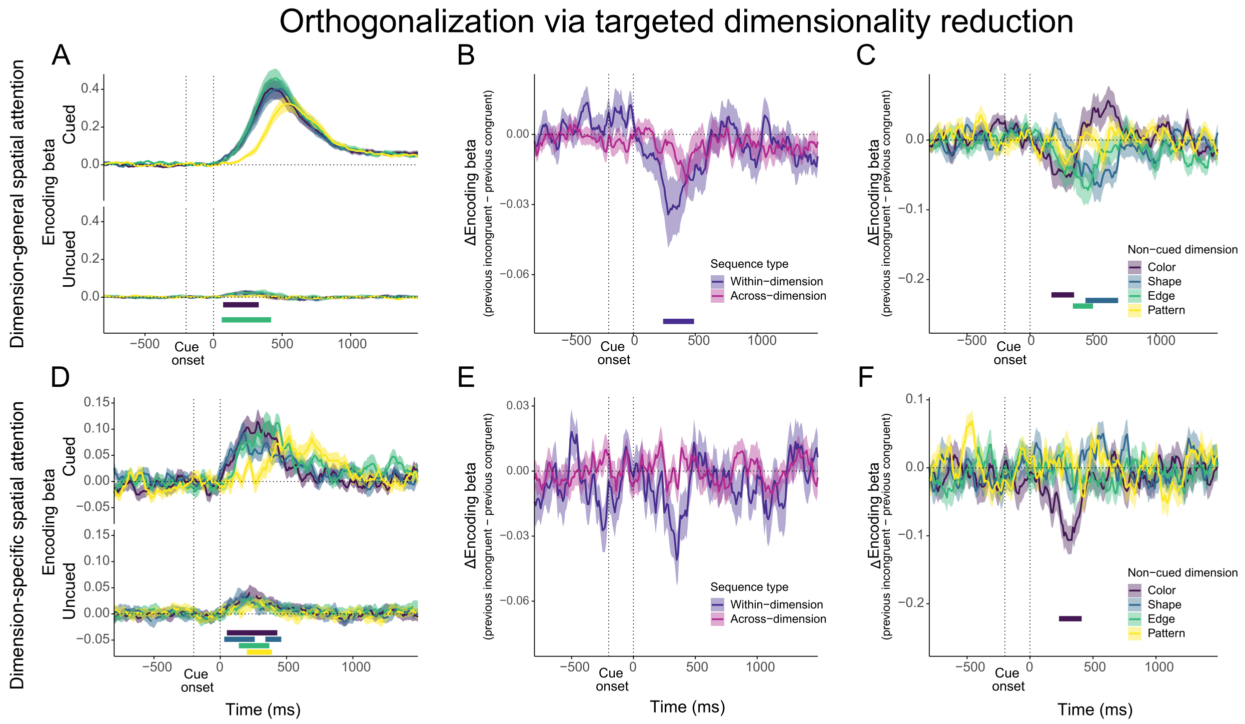


**Figure S3. Encoding model adaptation effects on distractor spatial attention. a,** Spatial orienting (i.e., dimension-general spatial attention) projections for cued (top) and non-cued (bottom) dimensions, split by dimensions. Cued-dimension attention rises strongly after stimulus onset; non-cued attention is weak and transient, with reliable clusters indicated by colored bars. **b,** Within- versus across-dimension adaptation effects on spatial orienting to distractors, computed as the difference in coding strength following incongruent versus congruent trials. Robust within-dimension suppression emerges after stimulus onset, with no reliable across-dimension effect. **c,** Within-dimension adaptation split by non-cued dimension, showing earlier and stronger effects for color and edge, weaker effects for shape, and absent effects for pattern. **d,** Feature coding (i.e., dimension-specific spatial attention) projections for cued (top) and non-cued (bottom) dimensions. A similar pattern is observed, with stronger and more sustained cued-dimension coding. Although smaller in magnitude, non-cued feature coding was more reliable and emerged earlier than non-cued spatial orienting, in line with the reactive curtailing of distractor processing. **e,** Within- versus across-dimension adaptation effects on distractor feature coding. Within-dimension suppression is reliable and mirrors the pattern in **b**. **f,** Within-dimension adaptation split by non-cued dimension for the feature coding axis, showing robust, fast, and selective adaptation for color. Shaded areas indicate ±1 SEM; vertical dashed lines mark cue and stimulus onsets; colored bars indicate significant clusters.

The feature coding axes (Fig. S3D–F) captured variance private to each dimension, potentially including visual feature processing, regardless of whether it served as the task-relevant target (cued dimension) or as a distractor (non-cued dimension). The analyses using this model confirmed that distractor features are encoded in parallel with target features, consistent with early perceptual processing preceding control (Fig. S3D). However, when we examined within-dimension adaptation, only the color dimension showed reliable modulation (Fig. S3EF). This suggests that most of the conflict adaptation captured by the decoding-RSA approach reflects adjustments along spatial orienting (i.e., the dimension-general spatial-attention axis). Alternatively, because color interference occurs early and with high potency, its rapid dynamics may have made it uniquely sensitive to feature-level adaptation.

Re-aligning trials to response onset further supported the interpretation that the modulation of spatial orienting towards distractors reflects attentional capture followed by suppression: attention toward the cued dimension increased further and peaked sharply at the time of response, whereas distractor-directed attention disappeared entirely prior to response execution (Fig. S4). Thus, the dimension-general spatial attention towards distractors reflected brief orienting responses rather than sustained engagement.

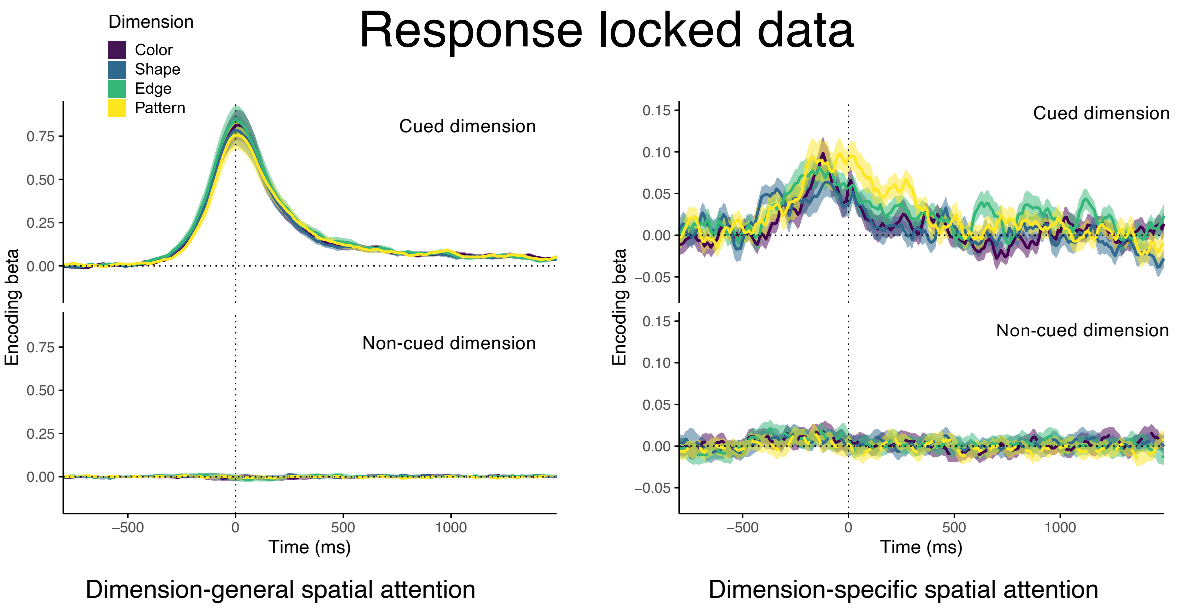


**Figure S4. Response-locked encoding model projections for cued and non-cued dimensions.** Projections onto the dimension-general attention axis (left – spatial orienting) and dimension-specific axis (right - feature coding) for cued (top) and non-cued (bottom) dimensions, shown separately for each task dimension (color-coded), time-locked to the response. Cued-dimension spatial orienting peaked at the response, whereas non-cued dimension orienting disappeared entirely before response execution, consistent with suppression preventing the stabilization of distractor-related spatial-attention signals. A similar dissociation is observed for the feature coding axis. Shaded areas indicate ±1 SEM; vertical dashed line marks response onset.
